# Developmentally-regulated splice remodeling shapes functional *CACNA1C* channel diversity during human neurodevelopment

**DOI:** 10.64898/2026.11.07.765057

**Authors:** Melis Çelik, Osama A. Arshad, Nicola A.L. Hall, S. Suheda Yasarbas, Mahnaz Nikpour, Lina Jonsson, Carl M. Sellgren, Mikael Landén, Antonios Pantazis, Paul J. Harrison, Arne W. Mould, Erik Smedler

## Abstract

*CACNA1C* encodes the pore-forming subunit of the Ca_V_1.2 L-type voltage-gated calcium channel, a major regulator of neuronal calcium signaling and one of the strongest genetic risk loci for bipolar disorder and other psychiatric disorders. Although extensive alternative splicing of *CACNA1C* has been described in the adult human brain, how alternative splicing of *CACNA1C* is regulated during early human neurodevelopment remains largely unknown. Here, we used long-read RNA sequencing to characterize full-length *CACNA1C* transcripts during differentiation of human induced pluripotent stem cells, from healthy controls and patients with bipolar disorder homozygous for either the A or G allele at rs1006737, into cortical glutamatergic neurons. We identified 60 putative isoforms, including ten annotated transcripts and 23 previously reported in adult human brain. We also demonstrate that *CACNA1C* undergoes developmentally-regulated remodeling of isoform composition during neuronal differentiation. This remodeling was characterized by an exon-switching program involving mutually exclusive exon pairs and alternative splice site usage, including prominent developmental transitions between exon pairs 8/8a, 21/22, and 31/32. Neither rs1006737 genotype, diagnosis, sex nor age significantly affected splicing profiles. The developmental splice trajectories were recapitulated in dorsal forebrain organoids, showing that *CACNA1C* splice remodeling is conserved across independent human neural differentiation models. Electrophysiological characterization of the two most abundant neuronal isoforms found that exon 31/32 switching alters channel inactivation properties. Together, our findings demonstrate that developmentally-regulated alternative splicing of *CACNA1C* transcripts contributes to the functional diversity of Ca_V_1.2 channels expressed during human neurodevelopment.

## Introduction

*CACNA1C* encodes the pore-forming α_1C_ subunit of the Ca_V_1.2 L-type voltage-gated calcium-selective channel, a major mediator of calcium influx following membrane depolarization in excitable cells. Ca_V_1.2 channels regulate diverse aspects of neuronal development and function, including neuronal differentiation, dendritic maturation, activity-dependent transcription, and synaptic plasticity throughout the developing and adult brain ^1–6^. In parallel, genome-wide association studies (GWAS) have consistently identified single-nucleotide polymorphisms (SNPs) in *CACNA1C* as one of the strongest common genetic risk loci for bipolar disorder (BD) and other major psychiatric disorders, including schizophrenia and major depression ^7–11^. Despite this strong genetic and neurobiological evidence, the molecular mechanisms through which *CACNA1C* contributes to human brain development and psychiatric illness remain incompletely understood ^12^.

A major source of Ca_V_1.2 functional diversity is extensive alternative splicing of *CACNA1C*. Alternative exon usage can modify channel activation and inactivation kinetics, calcium-dependent regulation, subcellular localization and pharmacological sensitivity, as exemplified by Timothy syndrome ^13–15^. Several alternatively spliced exons have been implicated in tissue-specific and developmental regulation of channel function, particularly mutually exclusive exon pairs that alter voltage-sensing and intracellular regulatory domains ^16, 17^. Although individual alternatively spliced exons have been characterized, how combinations of exons are coordinated across full-length transcripts during neuronal maturation is largely unknown. Conventional short-read RNA sequencing incompletely reconstructs transcript structures and cannot reliably resolve complex combinations of mutually exclusive exons and alternative splice-site usage within individual *CACNA1C* transcripts ^18^. Long-read sequencing technologies now enable direct characterization of full-length transcripts, providing an opportunity to define regulation of alternative splicing with substantially greater precision ^19^. Notably, recent long-read sequencing studies of adult human postmortem tissues have revealed a remarkable diversity of full-length transcripts expressed by *CACNA1C* ^20–22^.

More broadly, ion channels can undergo developmentally regulated changes in isoform expression that alter their functional properties as neurons mature ^23^. Defining such developmental changes may be particularly important for neuropsychiatric risk genes such as *CACNA1C*, since developmental variation in splice isoform composition could influence how genetic or illness-associated effects are interpreted. However, previous studies of *CACNA1C* transcript diversity have focused largely on mature human brain tissue, leaving the developmental regulation of *CACNA1C* splicing poorly understood. It therefore remains unknown how *CACNA1C* splice isoform composition changes during early human neurodevelopment.

The *CACNA1C* variant rs1006737 (G/A), located within intron 3, has been repeatedly associated with BD in genome-wide association studies ^11, 24–27^. rs1006737 has also been reported to act as an expression quantitative trait locus (eQTL), with genotype-dependent differences in *CACNA1C* expression and the abundance of specific transcript classes ^28^. These findings raise the possibility that genetic variation at this locus may influence *CACNA1C* transcript composition in addition to overall expression, providing a rationale for investigating whether rs1006737 is associated with alternative splicing. However, whether rs1006737 is associated with relative splice isoform expression or differential exon usage, particularly during human neural development, remains unresolved.

Neuronal maturation is accompanied by coordinated developmental splicing programs orchestrated by RNA-binding proteins including PTBP, RBFOX, and NOVA family members ^29–31^. These temporally regulated programs drive transitions between immature and mature neuronal states and are essential for the functional maturation of numerous neuronal proteins, particularly ion channels ^29^. Developmental regulation of ion channel splicing provides a mechanism for progressively tuning cellular excitability, calcium influx, and activity-dependent signaling as neurons mature ^1, 6^. Given the central role of Ca_V_1.2 in neuronal physiology, developmental regulation of *CACNA1C* splicing may therefore represent an important mechanism for generating functionally distinct Ca_V_1.2 isoforms during human neurodevelopment; potentially of importance in the pathophysiology of neuropsychiatric disorders.

Here, we tested the hypothesis that *CACNA1C* undergoes developmentally-regulated splice remodeling during human neurogenesis, generating functionally altered Ca_V_1.2 channels. We also investigated whether this remodeling was associated with BD diagnosis, rs1006737 genotype, sex or age. To address this, we combined PCR-targeted long-read RNA sequencing with human iPSC-derived neural differentiation, dorsal forebrain organoids, and electrophysiological characterization of abundant isoforms. We identified a developmentally-regulated exon-switching program involving mutually exclusive exons and alternative splice-site usage, demonstrated that these splice trajectories are conserved across independent human neural differentiation models, and showed that developmentally-regulated splice variants have different electrophysiological properties.

## Results

### Developmental modeling of human neural differentiation

To investigate developmental *CACNA1C* splice dynamics during human neurogenesis, fibroblasts from six individuals with BD and six healthy controls were reprogrammed into iPSCs using an integration-free synthetic RNA-based approach. Following quality control, iPSC lines from six BD patients and six controls, homozygous for either the A or G allele at rs1006737, were differentiated into neural progenitor cells (NPCs) and cortical glutamatergic neurons using a dual SMAD inhibition protocol ^32^. Samples were collected at the iPSC (DIV0, n=12 donors), NPC (DIV35, n=10 donors), and neuronal (DIV50, n=10 donors) stages for RNA isolation and subsequent bulk RNA sequencing and PCR-targeted long-read sequencing to resolve full-length *CACNA1C* isoforms (**Figure 1A** and **Supplementary Table 1** shows all iPSC lines and assays). Donor characteristics, including diagnosis status, rs1006737 genotype, age, sex, and BD subtype are provided in **Supplementary Table 2**.

**Figure 1.**
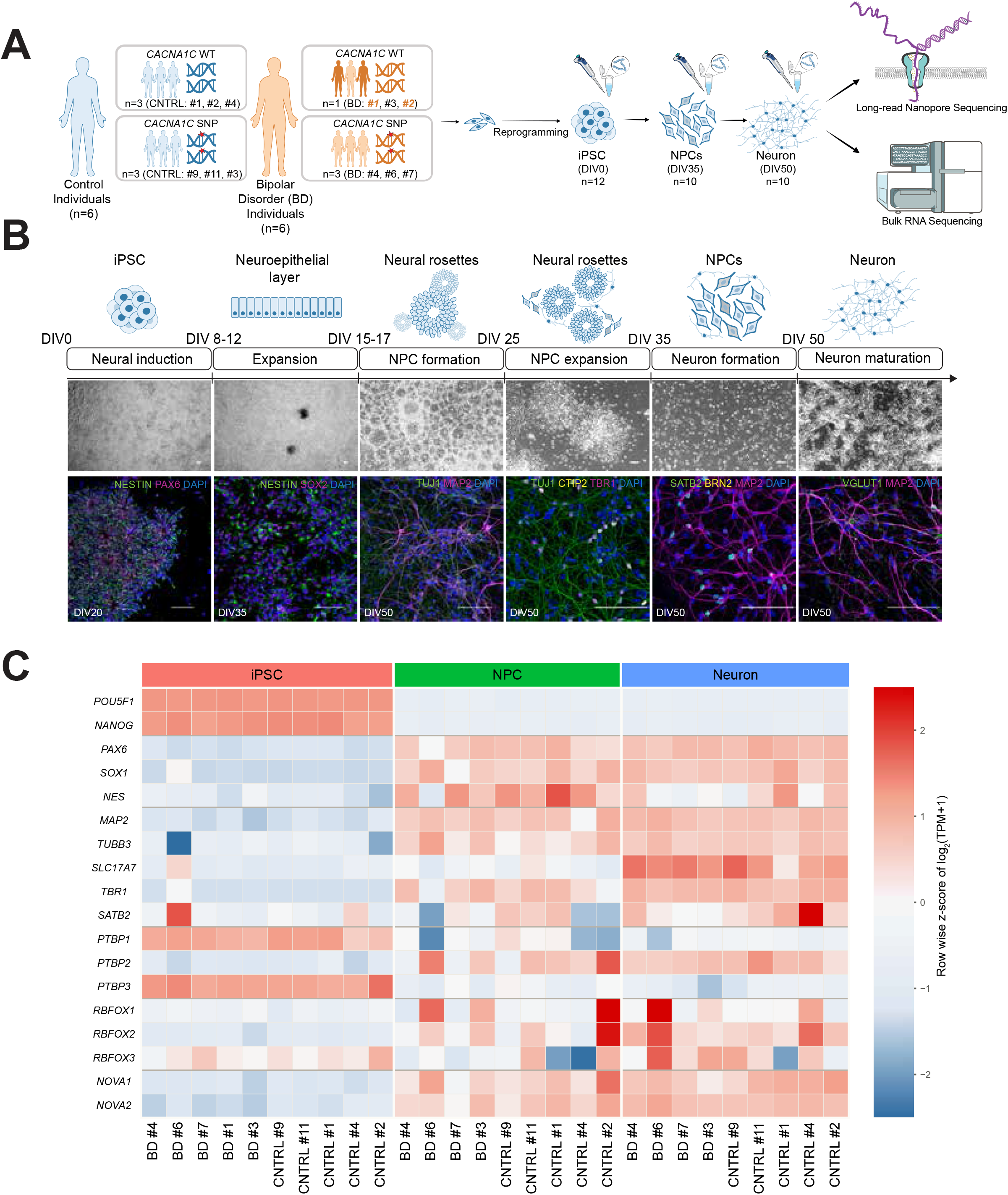
Splice isoform profiling across neural differentiation in patient and control iPSC-derived models using long-read sequencing. **(A)** Fibroblasts were isolated from six controls and six bipolar disorder patients, comprising six rs1006737 A homozygotes and six G homozygotes, and reprogrammed to iPSCs and differentiated to cortical neurons. Dark orange-labeled samples (two patients) were not included in Nanopore sequencing due to differentiation issues at NPC and neuron stages. RNA samples were collected over differentiation at three different stages, including iPSC (DIV0), NPC (DIV35), and Neuron (DIV50) for bulk RNA sequencing and Nanopore sequencing. For Nanopore sequencing, two excluded lines were included only in iPSC stages. **(B)** Schematic representation of dual SMAD inhibition protocol to differentiate from iPSCs to cortical neurons accompanied by representative brightfield images to represent critical points over differentiation from DIV0 to DIV50 Scale bar: 100 µm. The differentiation was confirmed by immunostaining to assess the expression of progenitor markers (NESTIN, SOX2, PAX6), neuronal markers (TUJ1, MAP2), and the glutamatergic marker vGLUT1 together with deep-layer (TBR1, CTIP2) and upper-layer (SATB2, BRN2) cortical markers. Scale bar: 100 µm. **(C)** Heatmap showing normalized bulk RNA-seq expression (TPM) of pluripotency markers (*POU5F1*, *NANOG*), progenitor markers (*PAX6*, *SOX1*, *NES*), cortical glutamatergic neuronal markers (*MAP2*, *TUBB3*, *SLC17A7*, *TBR1*, *SATB2*), and splicing regulators (*PTBP*, *RBFOX*, and *NOVA* family genes) across differentiation stages.

Successful neuronal differentiation was confirmed by immunostaining for stage-specific markers (**Figure 1B**). NPC identity and neural rosette formation were supported by the expression of NESTIN, PAX6, and SOX2, whereas glutamatergic neuronal identity was confirmed by MAP2, TUJ1, and vGLUT1 together with the expression of deep-layer (TBR1 and CTIP2) and upper-layer (SATB2 and BRN2) cortical markers. Bulk RNA sequencing further validated stage-specific transcriptional programs (**Figure 1C**), with enrichment of *POU5F1* and *NANOG* in iPSCs, *PAX6*, *SOX1*, and *NES* in NPCs, and *MAP2*, *TUBB3*, *SLC17A7*, *TBR1* and *SATB2* in neurons.

Splicing regulator genes (i.e., *PTBPs*, *RBFOX*, and *NOVA1/2*) showed stage-dependent expression profiles during differentiation from iPSCs to NPCs and neurons (**Figure 1C**). *PTBP1* and *PTBP3* were enriched in iPSC, and decreased in NPC and cortical neurons, whereas *RBFOX1/3* and *NOVA* family members were upregulated in neuronal states, consistent with the establishment of neuron-specific alternative splicing programs. *PTBP2* exhibited an intermediate pattern across differentiation stages, consistent with its reported role during neuronal maturation ^33^. These findings suggest a developmental transition from progenitor-associated PTBP1-dependent splicing networks toward mature neuronal RBFOX/NOVA regulatory programs.

### Long-read sequencing reveals developmentally-regulated alternative splicing of *CACNA1C* transcripts

PCR-targeted long-read RNA sequencing identified 60 potential *CACNA1C* isoforms across 32 samples from the neural differentiation experiment described above (12 iPSC, 10 NPC, and 10 neuronal samples, representing 12 donor lines, of which 10 were sampled across all three developmental stages). Of these, ten had Ensembl IDs (i.e., are previously annotated) and 23 have been reported in adult human brain ^20, 21^ (**Figure 2A**, **Supplementary Table 3**). Novel isoforms were required to meet predefined filtering criteria using the default parameters implemented in IsoLamp ^21^. The exon and intron structures of 60 *CACNA1C* isoforms are provided in BED format (**Supplementary Data 1**).

**Figure 2.**
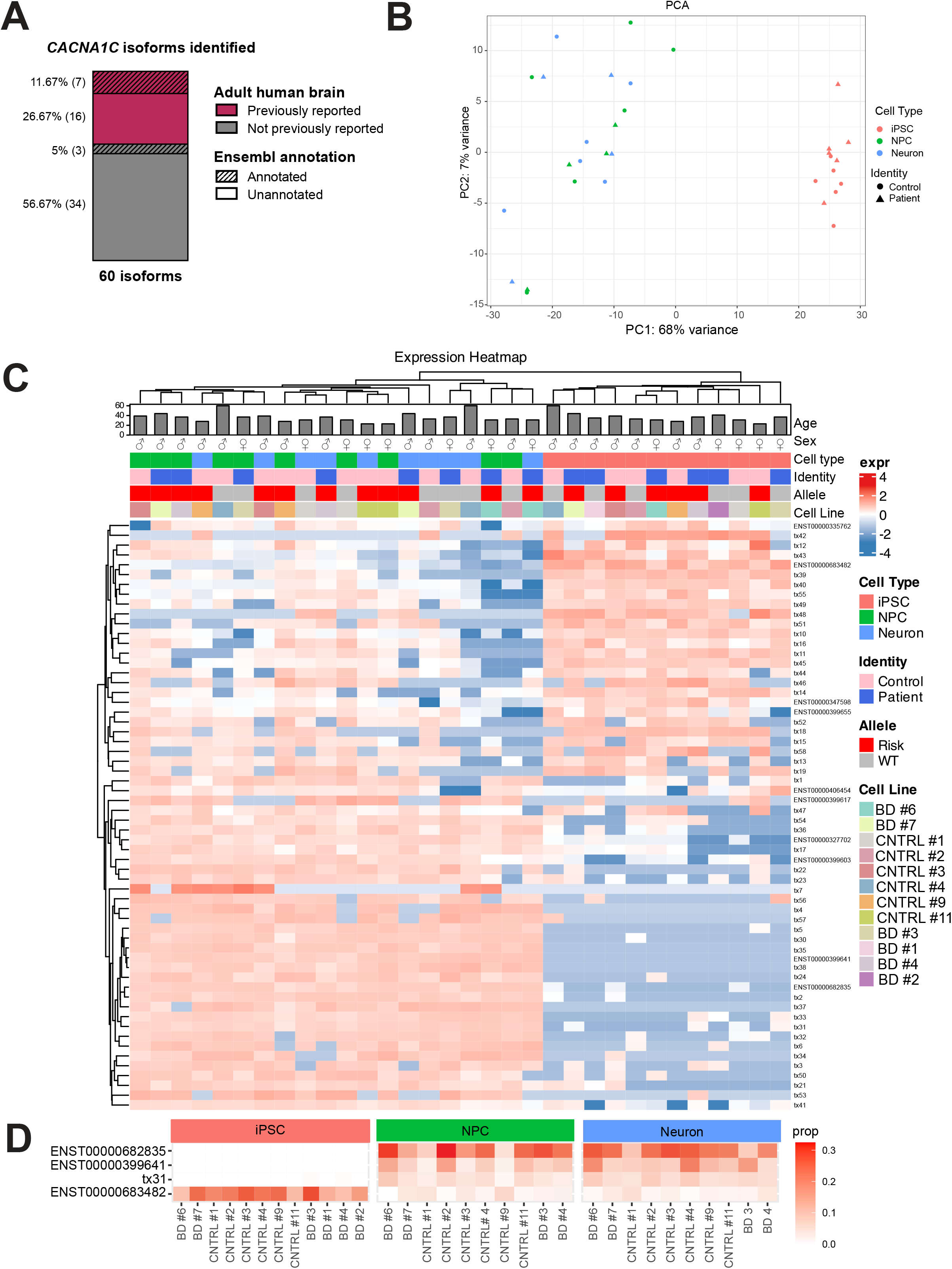
Isoform composition shifts markedly across differentiation stages. **(A)** Stacked column showing the distribution of 60 isoforms identified by Oxford Nanopore sequencing across differentiation stages. Pink indicates isoforms previously reported in adult human brain (n = 23), grey indicates ones not reported in the adult brain (n = 37). Hatching denotes Ensembl-annotated isoforms (n = 10), of which seven were previously reported in adult human brain. Percentages indicate the proportion of distinct isoforms in each category relative to 60 identified isoforms, with the corresponding number of isoforms shown in parentheses beside each percentage. **(B)** Principal component analysis (PCA) of Oxford Nanopore-derived *CACNA1C* isoform profiles demonstrating separation between iPSC and differentiated stages, while no segregation was observed based on diagnosis status. **(C)** Heatmap showing z-scaled log_2_ transformed expression of the 60 *CACNA1C* isoforms across all analyzed samples grouped by developmental stage, diagnosis status, rs1006737 genotype, sex, and donor age. **(D)** Relative expression proportions of the four most abundant *CACNA1C* isoforms across iPSC, NPC, and neuronal stages.

Principal component analysis revealed strong clustering according to developmental stage, with clear separation between iPSCs and neuronally differentiated samples (NPCs and glutamatergic neurons, **Figure 2B**). These findings indicate that *CACNA1C* splice remodeling follows highly structured developmental trajectories during human neural differentiation.

Heatmap visualization demonstrated stage-dependent isoform transitions across all samples, with no robust clustering according to diagnosis status, rs1006737 genotype, age, or sex (**Figure 2C**). Overall isoform composition was strongly associated with developmental stage in the balanced repeated-measures cohort of 30 samples from 10 donors (Hellinger–Euclidean PERMANOVA, R² = 0.671, pseudo-F = 27.54, Holm-adjusted *p* < 0.001). Developmental stage accounted for 67.1% of the multivariate variation in relative isoform composition, with permutations restricted within donor to account for repeated measurements across developmental stages (**Supplementary Figure 1**, **Supplementary Table 4**). The stage association was not accompanied by evidence of heterogeneous multivariate dispersion among stages (PERMDISP, *p* = 0.577). In separate stage-adjusted explanatory analyses, in which each donor’s three repeated samples were permuted together, *rs1006737* genotype (R² = 0.009, *p* = 0.716), diagnosis status (R² = 0.008, *p* = 0.773), sex (R² = 0.012, *p* = 0.392), and age (R² = 0.010, *p* = 0.544) explained little variation and were not statistically significant. All four associations remained nonsignificant after Holm correction. These findings were consistent with the stage-specific clustering observed in the isoform expression heatmap (**Figure 2C**). Post-hoc comparisons showed that the overall isoform composition differed between iPSC and NPCs (R² = 0.663, Holm-adjusted *p* = 0.006) and between iPSCs and neurons (R² = 0.710, Holm-adjusted *p* = 0.006), whereas there was no statistically detectable difference between NPCs and neurons (R² = 0.045, Holm-adjusted *p* = 0.318), which indicates that the overall developmental-stage association was primarily attributable to changes occurring between the iPSC and two other stages.

Relative isoform abundance analysis further revealed distinct dominant transcript isoforms before and after differentiation (**Figure 2D**, **Supplementary Figure 2**). The ten most abundant isoforms in iPSCs, NPCs and neurons are shown in **Figure 3** and **Supplementary Figure 3**. Isoform ENST00000683482 was most abundant in iPSCs whereas ENST00000682835 was most abundant in NPCs and neurons. Overall, eight of the ten most abundant isoforms differed between iPSCs and neurons. Of the 20 top isoforms seen in the iPSC-derived neurons, 15 have been reported in adult human brain tissue ^20, 21^ (**Supplementary Table 3**). Together, these results demonstrate extensive developmental remodeling of *CACNA1C* splice isoforms during human neurogenesis, and considerable but partial overlap between iPSC-derived neurons and adult brain.

**Figure 3.**
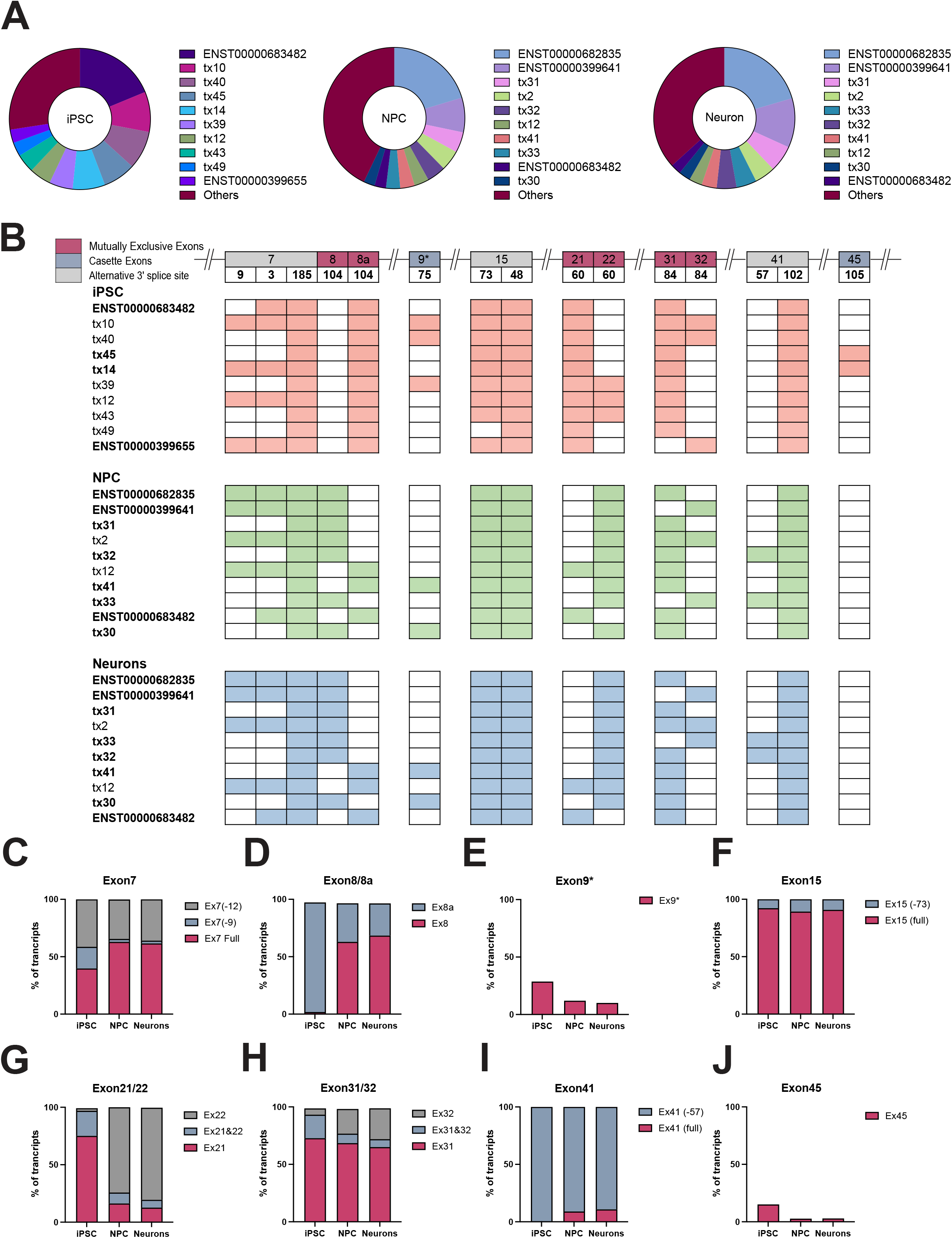
Developmental *CACNA1C* splice remodeling is driven by exon-specific alternative splicing patterns. **(A)** Proportion of transcript reads from the ten most abundant transcripts across three distinct developmental stages. **(B)** Exon composition of the top ten most abundant *CACNA1C* isoforms identified across iPSC, NPC, and neuronal stages, illustrating stage-dependent isoform transitions driven by alternative splicing of exons 7, 8/8a, 9*, 15, 21/22, 31/32, 41, and 45. For percentage contribution of each isoform to the transcript pool, see **Supplementary** Figure 3. Isoforms labelled in bold are predicted to give rise to a functional channel; isoforms containing truncated exon 15, exons 21 *and* 22, or exons 31 *and* 32, do not. **(C–J)** Quantification of mutually exclusive exon usage across differentiation stages for mutually exclusive exon pairs 8/8a **(D)**, 21/22 **(G)**, and 31/32 **(H)**. Quantification of additional stage-dependent alternative splicing events involving exons showing alternative 3’splice site (acceptor) usage in exons 7 **(C),** 15 **(F)**, 41 **(I)** and cassette exon usage involving exons, **(E)** 9* and 45 **(J)** Quantifications represent averaged exon usage across all lines (n=12 iPSC, n=10 NPC, and n=10 Neuron).

### Multiple exon-switching events characterize developmental *CACNA1C* splice remodeling

To define the structural basis of developmental splice remodeling, we next examined exon composition across the *CACNA1C* isoforms at the iPSC, NPC, and neuronal stages (**Supplementary Table 5)**. *CACNA1C* isoform composition showed a distinct profile at the iPSC stage compared with NPC and neuronal stages, which exhibited more similar isoform profiles (**Figure 3A**). Stage-dependent changes in isoform composition were associated with alternative splicing events involving mutually exclusive exons 8/8a, 21/22 and 31/32; cassette exons 9* and 45; and alternative 3’ splice-site (acceptor) usage in exons 7, 15 and 41 (**Figure 3B**, Supplementary Figure 3, Supplementary Table 6).

The mutually exclusive exon pair 8/8a exhibited a pronounced developmental transition, with exon 8a predominating in iPSCs and exon 8 becoming dominant following neural differentiation (**Figure 3D).** Both splice forms demonstrated significant developmental stage associations (Friedman tests, BH-adjusted *p* = 0.0012), with significant difference between iPSCs and both NPCs and neurons (paired Wilcoxon tests, BY-adjusted *p* = 0.0268). Similarly, exon pair 21/22 showed a developmental shift from exon 21 predominance in iPSCs to exon 22 predominance following differentiation stages (**Figure 3G**). Both single exon configurations exhibited significant developmental stage associations (BH-adjusted *p* = 0.0012), with significant differences between iPSCs and both differentiated stages (BY-adjusted *p* = 0.0268). Exon pair 31/32 underwent a more gradual transition, with exon 31 remaining predominant throughout differentiation while the relative contribution of exon 32 increased from iPSCs to NPCs and neurons (**Figure 3H**). The exon 32 showed a significant developmental stage association (BH-adjusted *p* = 0.0012), with significant differences between iPSCs and both differentiated stages (BY-adjusted *p* = 0.0268). In addition, total exon 31 inclusion showed a significant developmental stage association (BH-adjusted *p* = 0.003).

Although exons 21/22 and 31/32 are annotated as mutually exclusive, isoforms containing co-inclusion of these exons were also detected. However, they had low abundance and are known to result in a protein unable to function as a channel ^34^. Whilst these considerations require caution as to their significance, isoforms containing both exons 21 and 22 have been identified previously ^20^ and reported to have a dominant negative effect by competing for the beta subunit of the channel ^34^.

Additional stage-dependent splice remodeling was observed across several regions. Exon 7 exhibited alternative 3′ splice-site (acceptor) usage, including isoforms with 9- or 12-bp deletions; the 9-bp deletion was enriched in iPSCs, whereas full-length exon 7 predominated in NPCs and neurons (Friedman tests, BH-adjusted *p* = 0.0012 for both features, **Figure 3C**). The 12-bp deletion showed no significant stage association. Exon 15 similarly displayed alternative 3′ splice-site (acceptor) usage, generating truncated variants slightly more abundant in mature stages, with an overall stage association (BH-adjusted *p* = 0.0081, **Figure 3F**). Differential usage of exon 41 was also observed, with inclusion of the extended exon detected in NPCs and neurons but negligible in iPSCs (BH-adjusted *p* = 0.0012, **Figure 3I**). Exon 45 usage was largely enriched in the iPSC stage (BH-adjusted *p* = 0.0012, **Figure 3J**). Likewise, the other cassette exon, 9*, was more abundant in iPSCs (BH-adjusted *p* = 0.0038, **Figure 3E).**

To determine whether *CACNA1C* exon splicing patterns differed independently of developmental stage, exon-splicing contributions at sample-level (per line and stage) were compared between patients and controls, rs1006737 A and G homozygotes, and sex, separately at the three developmental stages (**Supplementary Table 7**). No individual splicing feature differed significantly between patients and controls at any developmental stage after multiple-testing correction (all FDR > 0.05). Similarly, no significant differences were detected between rs1006737 A and G homozygotes or between females and males at any of the three developmental stages (all FDR > 0.05). Overall, no diagnosis-, rs1006737 genotype-, or sex-associated alterations in the examined *CACNA1C* exon-splicing features were detected when samples were compared at equivalent developmental stages.

### Developmental *CACNA1C* splice remodeling is recapitulated in dorsal forebrain organoids

To investigate whether these developmental splice trajectories in two-dimensional cultures were recapitulated in a more physiologically relevant three-dimensional system, we next examined *CACNA1C* splicing in dorsal forebrain organoids ^35^ derived from human embryonic stem cells. To capture progressive maturation states, dorsal forebrain organoids were analyzed at DIV50, DIV150, and DIV180. Dorsal forebrain organoid identity and maturation were confirmed using stage-specific and regional markers (**Figure 4A**; **Supplementary Figure 4**). Splice dynamics were investigated using targeted qPCR assays designed to quantify key mutually exclusive exon pairs (**Figure 4B**).

**Figure 4.**
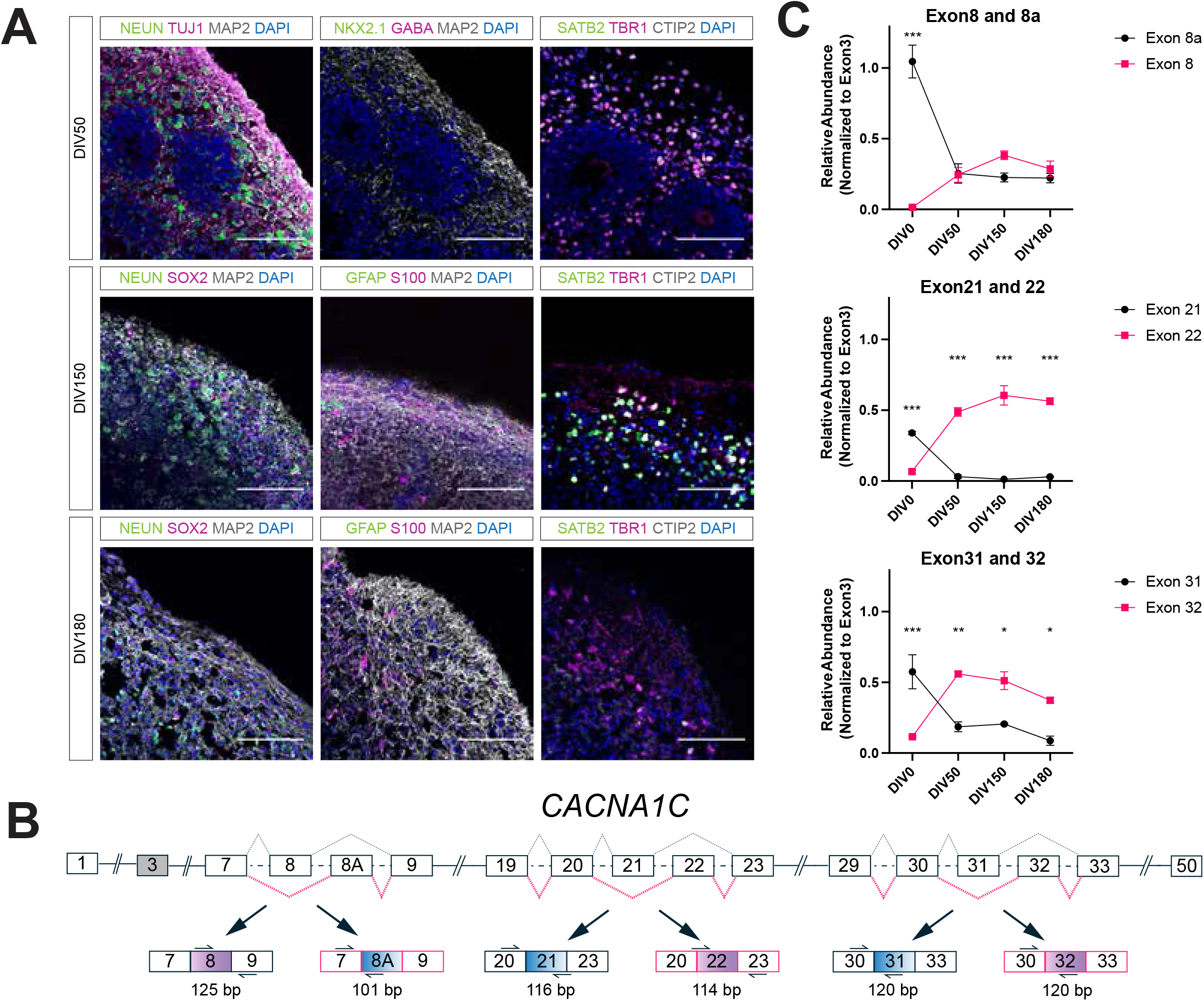
Developmental *CACNA1C* splice dynamics are recapitulated in dorsal forebrain organoids. **(A)** Representative validation of dorsal forebrain organoid identity and maturation at DIV50, DIV150, and DIV180 using cortical layer markers (TBR1, CTIP2, SATB2), ventral markers (NKX2.1, GABA), neuronal maturation markers (TUJ1, NEUN, MAP2), progenitor marker SOX2, and glial markers (S100, GFAP). Representative immunostaining images were obtained from 10 µm sections of 1–3 independently processed organoids per developmental time point. Scale bar, 100 µm. **(B)** Experimental overview of RT-qPCR assays targeting mutually exclusive *CACNA1C* exon pairs (8/8a, 21/22, and 31/32) during dorsal forebrain organoid maturation. Arrows indicate exon-specific primer binding sites used for selective amplification of targeted splice variants. Relative exon usage was normalized to constitutively expressed exon 3 (marked gray) across developmental stages. **(C)** RT-qPCR quantification of mutually exclusive exon usage during organoid maturation. Expression levels were normalized to *GAPDH* and constitutive exon 3 to assess the relative contribution of targeted exons across developmental stages. Data represent mean ± SEM (n = 3 biologically independent organoids per developmental time point, with each sample measured in technical duplicate.). Statistical significance was evaluated using two-way mixed ANOVA, with exon identity as the repeated-measures factor, followed by Šídák’s multiple-comparisons test comparing exon usage within each developmental stage. Significant time x exon interactions were observed for exon 8/8a (F(3,8) = 38.86, *p* < 0.0001), exon 21/22 (F(3,8) = 92.36, *p* < 0.0001), and exon 31/32 (F(3,8) = 29.40, *p* = 0.0001). Significance symbols indicate Šídák-adjusted comparisons between exons at each developmental time point: \**p* < 0.05, \*\**p* < 0.01, \*\*\**p* < 0.001.

Consistent with observations from the two-dimensional neural differentiation cultures, dorsal forebrain organoids exhibited developmental *CACNA1C* splice remodeling, suggesting that similar splicing dynamics occur across structurally distinct two- and three-dimensional human neural differentiation systems (**Figure 4C**).

### Developmentally-regulated *CACNA1C* exons modulate channel inactivation

To investigate whether developmental splice remodeling alters Ca_V_1.2 channel function, we examined the two most abundant iPSC-neuron isoforms, which differ in the inclusion of either exon 31 or exon 32 in the S3–S4 linker of domain IV: ENST00000682835 (Ca_V_1.2-241, top iPSC-neuron isoform) and ENST00000399641 (Ca_V_1.2-217, top adult brain isoform^20^, **Figure 5A**, **Figure 6**). Electrophysiological recordings of heterologously-expressed, neuronal-like Ca_V_1.2 channel complexes (α_1C_/α_2_δ-1/β_4_) revealed similar voltage-dependent activation properties in Ca_V_1.2-217 and Ca_V_1.2-241 (**Figure 5B-D, Supplementary Table 8**).

**Figure 5.**
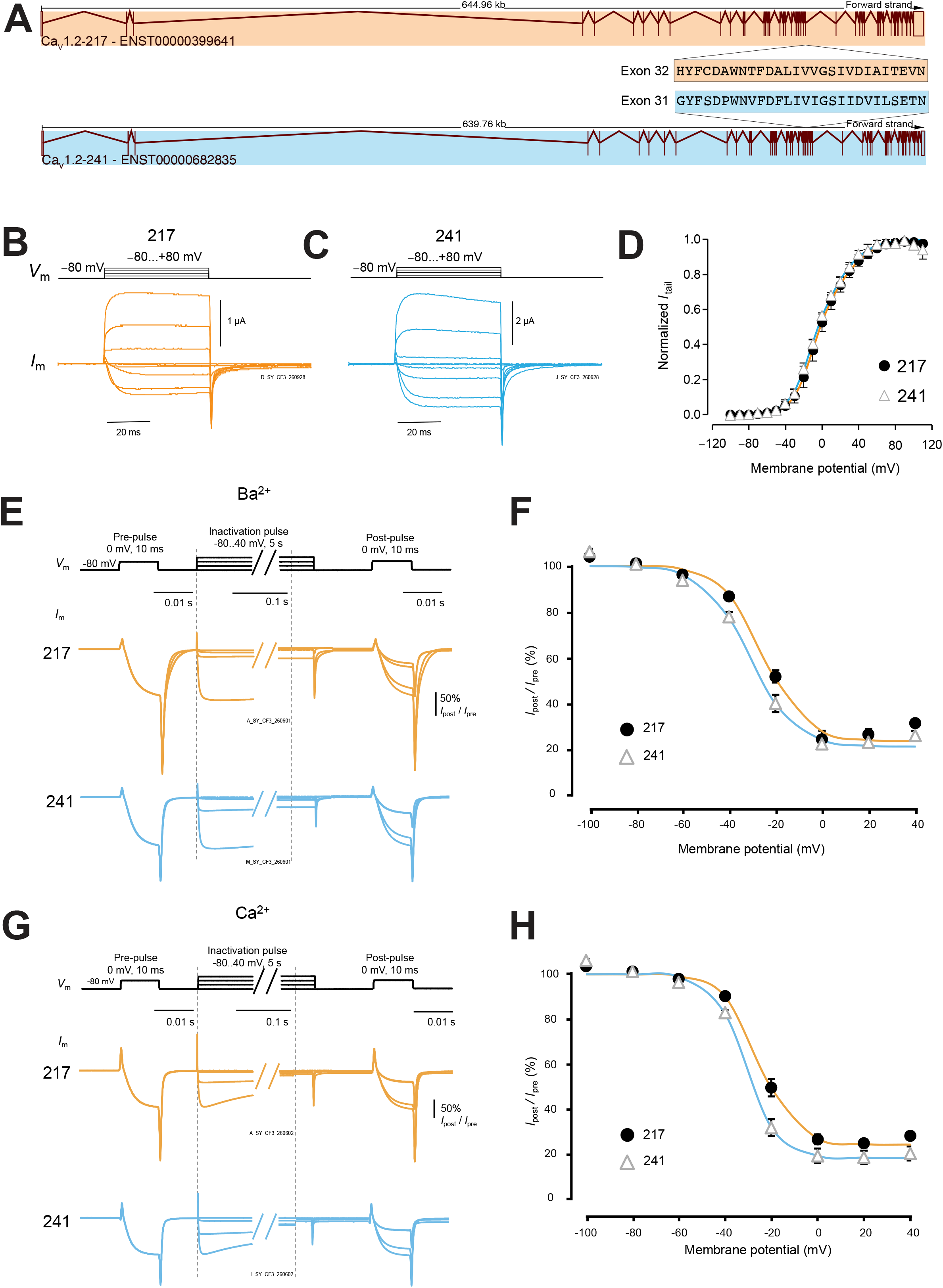
Exon 31/32 usage modulates Ca_V_1.2-channel inactivation. (A) Transcript structure of *CACNA1C*-217 (ENST00000399641, referred here as ‘217’) and *CACNA1C*-241 (ENST00000682835, referred here as ‘241’) showing exon-intron organization and transcript orientation, differing only in exon 31/32 usage (*viz.*, 241 contains exon 31, 217 exon 32). **(B)** Membrane potential protocol (*V*_m_) for channel activation. Representative current traces (*I*_m_) for 217 elicited by 50-ms test pulses. **(C)** As in panel B, showing currents from 241. **(D)** Voltage dependence of normalized tail currents (*I*_tail_) for 217 (black filled circles) and 241 (open grey triangles). Curves represent fits to the sum of two Boltzmann functions (parameters in **Supplementary Table 8**). **(E)** Membrane potential protocol (*V*_m_) for channel inactivation. A 5-s inactivating pulse to the indicated potentials was flanked by 10-ms pre- and post-pulses to 0 mV to assess channel availability before and after the inactivating steps. Dashed lines mark changes in the time scale. Representative Ba² current traces (*I*_m_) are shown below. Note that the traces were normalized to the amplitude of the pre-pulse current; raw traces are shown in **Supplementary** Figure 5. **(F)** The voltage dependence of inactivation in Ba² for 217 (filled black circles) and 241 (open grey triangles). Curves represent fits to a Boltzmann function (parameters in **Supplementary Table 8**). **(G, H)** As in panels E and F, respectively, for inactivation experiments in calcium. Raw current traces in **Supplementary** Figure 5 and Boltzmann parameters in **Supplementary Table 8**. Errors are SEM.

**Figure 6.**
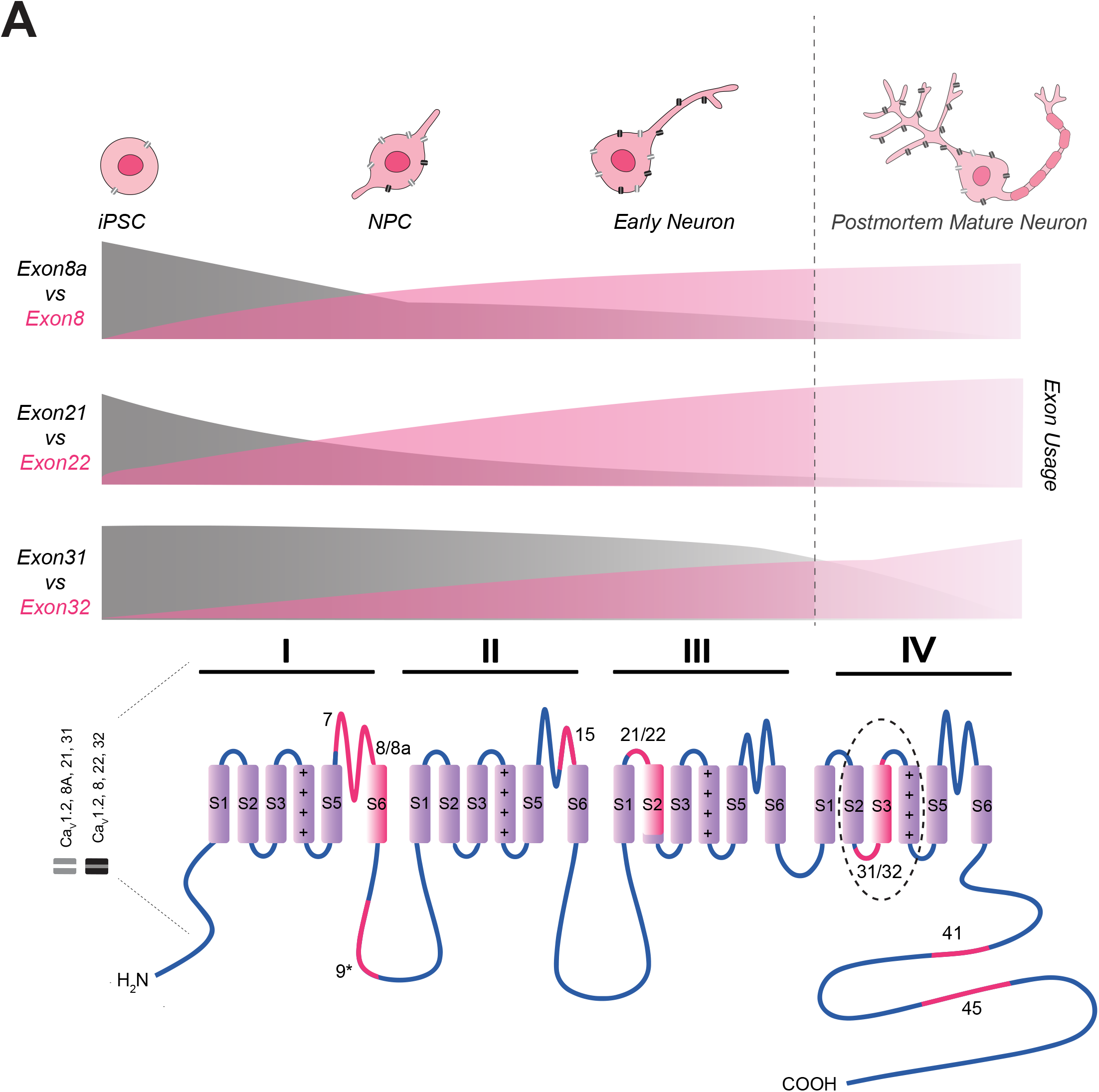
Graphical overview of *CACNA1C* splicing regulated during human neurodevelopment. Graphical model summarizing stage-dependent *CACNA1C* isoform remodeling with the contribution of mutually exclusive exons. Gradients illustrate qualitative patterns and are not proportional to measured abundances (inspiration from ^23^).

We also studied the voltage dependence of inactivation, in external solution containing barium or calcium: the former precluded calcium-dependent inactivation (CDI), showing only voltage-dependent inactivation (VDI); while the latter allowed the study of both VDI and CDI ^36, 37^, with the physiological charge carrier. ^38–41^ We found that in Ca_V_1.2-217, VDI was inhibited relative to Ca_V_1.2-241, with a voltage dependence shifted to more positive voltages by ∼6 mV (**Figure 5E-F, Supplementary Figure 5A, Supplementary Table 8**). This difference was also observed in the presence of calcium / CDI, highlighting its physiological relevance (**Figure 5G-H, Supplementary Figure 5B, Supplementary Table 8**). Together, these findings indicate that a developmental transition, from Ca_V_1.2-241 in immature neurons that iPSC-neurons represent to Ca_V_1.2-217 in fully mature neurons, would be associated with altered inactivation, linking developmental *CACNA1C* splice remodeling to changes in Ca_V_1.2 channel availability.

## Discussion

Alternative splicing is a major source of proteomic diversity in the nervous system, yet how full-length splice isoforms are developmentally regulated in genes central to neuronal signaling remains poorly understood. Here, using PCR-targeted long-read sequencing of full-length *CACNA1C* transcripts, we demonstrate that *CACNA1C* undergoes developmentally-regulated splice remodeling during human neurodevelopment. Rather than exhibiting a static repertoire of splice isoforms, *CACNA1C* showed developmental variation in exon usage during neuronal differentiation involving multiple alternatively spliced regions. These developmental trajectories were reproducible across both two-dimensional neural differentiation cultures and dorsal forebrain organoids and were accompanied by functional differences in Ca_V_1.2 channel inactivation, indicating that developmental splice remodeling contributes to the emergence of functionally diverse channel states during neuronal maturation.

Previous studies have established that *CACNA1C* is extensively alternatively spliced in the adult human brain and that individual splice events influence channel biophysics and illness phenotypes, including Timothy syndrome ^13, 14, 20^. However, these studies have largely focused on mature tissues and therefore provide limited insight into how transcript diversity emerges during development. The developmental focus is particularly relevant given accumulating evidence that genetic risk for psychiatric disorders converges on early brain development, with many risk-associated genes exhibiting peak expression or functional importance during fetal and early postnatal stages ^42–44^. Our findings extend previous observations by placing *CACNA1C* splice diversity into a developmental framework. This developmental perspective may be particularly important for neuropsychiatric risk genes, whose molecular and functional states can differ substantially between immature and mature neural cells ^23^. Rather than expressing a fixed transcript repertoire, *CACNA1C* expression undergoes progressive splice remodeling throughout neuronal differentiation, with distinct exon regions following different developmental trajectories. The most abundant isoform in adult human brain ^20^ is only second most abundant in iPSC-neurons here, indicating that further splice remodeling occurs during development to fully mature neurons.

One of the most striking observations was the regulation of multiple alternatively spliced regions. The mutually exclusive exon pairs 8/8a, 21/22, and 31/32 each exhibited characteristic developmental transitions, although with distinct temporal profiles (**Figure 3**). To our knowledge, this is also the first demonstration of developmental regulation of the mutually exclusive exon 21/22 pair during human neuronal differentiation. Exon 8/8a switching occurred early during neural differentiation, whereas exon 31/32 remodeling followed a more gradual pattern in 2D cultures and remained evident during prolonged organoid maturation (up until 180 days *in vitro*). These observations suggest that developmental remodeling of *CACNA1C* is not driven by independent regulation of individual exons but instead reflects a coordinated developmental splice program that progressively reshapes channel composition during neuronal maturation. Such sequential remodeling may allow Ca_V_1.2 channels to acquire altered physiological properties as developing neurons transition from proliferative progenitors to mature excitatory neurons.

Our observations are consistent with the broader concept that neuronal maturation is accompanied by coordinated developmental splicing programs. During neurodevelopment, transitions in the expression of RNA-binding proteins, including PTBP, RBFOX and NOVA family members, orchestrate large-scale changes in exon selection that are required for neuronal maturation rather than simply neuronal identity ^29, 31, 33^. Consistent with these established programs, we observed progressive downregulation of *PTBP* family members together with increased expression of *RBFOX* and *NOVA* genes across differentiation (**Figure 1C**). Although our data do not identify the upstream regulators responsible for *CACNA1C* splice remodeling, they place *CACNA1C* within the developmental splicing landscape that accompanies neuronal maturation. Future studies integrating transcriptome-wide splicing analyses with RNA-binding protein occupancy will be required to determine the regulatory mechanisms governing exon selection within the *CACNA1C* locus.

The functional significance of developmental splice remodeling is supported by our electrophysiological analyses. Developmentally regulated exon 31/32 switching altered voltage-dependent inactivation of Ca_V_1.2, while voltage-dependent activation was largely unchanged, demonstrating that this developmental splice event generates channels with distinct gating properties (**Figure 5**). Interestingly, these findings differ from previous heterologous expression studies reporting little or no difference in activation or inactivation between exon 31- and exon 32-containing Ca_V_1.2 channels ^17, 45^. One possible explanation is that the functional effect of exon 31/32 depends on the broader splice background in which these mutually exclusive exons occur. Notably, the isoforms examined in previous studies contained exon 8a rather than exon 8, which is present in the isoforms examined here, and exon 8/8a usage is itself known to influence voltage-dependent inactivation ^15^. Previous studies have also demonstrated interactions between *CACNA1C* splice events in determining channel properties ^17, 45^, suggesting that the functional consequences of exon 31/32 switching may be context dependent.

The location of exons 31/32 within the S3–S4 linker of voltage-sensor domain IV (VSD IV) provides a potential mechanistic basis for the selective effect on inactivation. The four Ca_V_1.2 voltage-sensor domains have distinct functional properties, and previous work indicates that activation of VSDs I–III provides the major energetic contribution to pore opening ^46, 47^. Consistent with this, exon 31/32 switching did not measurably alter voltage-dependent channel activation in our experiments (**Figure 5B–D; Supplementary Table 8**). By contrast, VSD IV activates more slowly than channel opening and has been implicated in slower voltage-dependent processes, including inactivation^46^, providing a plausible mechanistic explanation for the different inactivation properties of Ca_V_1.2-217 and Ca_V_1.2-241 (**Figure 5E–H**; **Supplementary Table 8**). Developmental alternative splicing also occurs within the corresponding repeat IV S3–S4 linker of the skeletal-muscle L-type channel Ca_V_1.1, where exon inclusion markedly modifies channel gating ^48^, further supporting this region as a functionally sensitive target of developmental splice regulation.

By altering the amplitude of Ca_V_1.2-mediated calcium influx, alternative splice isoforms are expected to modify excitation-transcription coupling and activity-dependent gene expression, processes that themselves contribute to neuronal maturation and circuit refinement^1, 6^. We therefore speculate that *CACNA1C* splicing is embedded within reciprocal interactions between intrinsic developmental splicing programs and activity-dependent signaling, forming feed-forward mechanisms that reinforce developmental progression. Rather than acting as independent processes, intrinsic developmental splicing programs and activity-dependent calcium signaling are likely to cooperate during neuronal maturation. *CACNA1C* may occupy a unique position at the interface between these processes.

Validation of the major developmental splice transitions in dorsal forebrain organoids strengthens the biological relevance of our findings. Although both differentiation systems model early human cortical development, they differ substantially in cellular organization and maturation. The observation that similar developmental exon-switching trajectories occurred in both systems argues that *CACNA1C* splice remodeling represents a robust feature of human neurodevelopment rather than an artifact of a particular differentiation protocol. At the same time, the sustained predominance of exon 32 during prolonged organoid maturation suggests that remodeling of *CACNA1C* splice isoforms extends beyond the developmental window captured by our two-dimensional neuronal cultures. This observation is also consistent with previous studies showing that cerebral organoids primarily recapitulate prenatal and early postnatal stages of cortical development rather than adult cortex^49^. Future studies using recently developed longer-term organoid culture systems may determine whether *CACNA1C* splice remodeling continues throughout later stages of human cortical maturation^50^. The similarities and differences from the *CACNA1C* isoform repertoire identified in the adult human brain have already been mentioned. Whilst the similarities attest to the fidelity of the iPSC-based approaches used here, the fact that many isoforms are not shared between the two datasets may reflect maturational stage, as well as a contribution from *CACNA1C* expressed in brain cells other than glutamatergic neurons (e.g. astrocytes), given that the latter studies used bulk tissue.

Our study has several limitations. First, the analyses were restricted to early stages of human cortical differentiation and targeted sequencing of a single gene. Consequently, although the observed splice transitions coincide with established neuronal developmental splicing programs, the upstream regulatory mechanisms remain unresolved. Second, stem-cell-derived neuronal cultures and organoid model early human neurodevelopment but do not fully recapitulate mature cortical circuitry^51^ nor expression of *CACNA1C* in all cell types. Third, the nature of PCR means that the sequencing results are only semi-quantitative and many of the isoforms were detected at low levels, requiring caution when interpreting their significance. Finally, while the present cohort included bipolar disorder patients and healthy controls, as well as rs1006737 A and G homozygotes, no associations of *CACNA1C* splicing with either diagnosis or rs1006737 genotype status were detected, indicating that developmental stage was the dominant determinant of transcript composition within this dataset. Much larger cohorts would be required to determine whether robust diagnosis- or genotype-specific alterations exist superimposed upon the developmental splicing program described here.

In conclusion, our findings establish *CACNA1C* as a dynamically regulated component of human neuronal development. We demonstrate that progressive remodeling of full-length splice isoforms accompanies neuronal maturation, is reproducible across independent human developmental models, and generates functionally different Ca_V_1.2 channel states. Together, our findings suggest that developmental remodeling of ion channel splice isoforms represents a mechanism through which intrinsic developmental programs and activity-dependent signaling become integrated during human cortical development.

## Methods

### Skin Biopsy and Fibroblast Derivation Procedure

A dermal biopsy was collected by a physician investigator^52^. After applying a local anesthetic patch, a 4 mm punch tool was used to extract a skin biopsy from the nondominant forearm. The biopsy samples were then placed in a transport medium of DMEM (Thermo Fisher Scientific, Cat. 11995065) supplemented with 1% Penicillin/Streptomycin (Gibco, Cat. 15140122) and 1% Glutamax (Gibco, Cat. 35050-038). Upon arrival at the laboratory, the samples were washed twice with PBS containing 10% Penicillin-Streptomycin-Glutamax. Two 60 mm tissue culture-treated plates were prepared (one for cutting the biopsy and the other for plating the tissue fragments). A scalpel was used to cut the biopsy into the smallest possible pieces, which were then arranged dermis-side down on a pre-marked grid in the plating dish. The dish was incubated at 37°C, with ambient oxygen and 5% CO_2_, under sterile conditions in an incubator for 15 minutes to allow adherence of the tissue. DMEM supplemented with 10% FBS and 1% Penicillin-Streptomycin-Glutamax was carefully added over the pieces of skin. The medium was changed twice a week. When the fibroblast started to emerge and covered the well, they were passaged by TrypLE™ Express Enzyme (1X), phenol red (ThermoFisher, Cat. 12605010) after being washed once with PBS. Gathered fibroblast cells were cultured in DMEM supplemented with 10% FBS and 1% Penicillin-Streptomycin-Glutamax in T75 plates (Sarstedt, Cat. 83.3911.002). The cells were tested for mycoplasma.

### Human Stem Cell Lines and Ethical Considerations

This study used human iPSCs and human embryonic stem cells (hESCs). The study was conducted under the ethical approval of the St. Göran Bipolar Project ^53^, and all experiments involving human material followed the criteria for ethical approval (2015/618-32). Reprogramming of cells was performed using integration-free synthetic RNA-based reprogramming by BrainXell. All cell lines were cultured at 37°C with 5% CO_2_, under sterile conditions in Forma Steri-Cycle CO_2_ Incubator Model 381 (Thermo Scientific). All cell treatments were performed in a sterile flow hood (n-Safe Pro One 1200, Nordic Labtech) using sterile, autoclaved instruments and media. iPSCs were screened for stem cell properties by evaluating their ability to differentiate and by confirming the expression of pluripotent stem cell markers *in vitro*. The human embryonic stem cells used in this study to generate dorsal forebrain organoids were obtained from the commercially available WA01/H1 hESC line, which has an XY 46 chromosome karyotype (WAe001-A, RRID:CVCL_9771; WiCell Research Institute).

### Karyotyping

The chromosomal stability of all healthy control and patient iPSC lines was assessed by classical karyotyping. All iPSC lines were cultured in T25 flasks coated with 5 μg/cm² Biolaminin LN521 (Biolamina, Cat.LN521-05) in mTeSR+ medium (STEMCELL Technologies, Cat. 100-0276) until they reached 70–80% confluency. To induce metaphase arrest, cells were incubated with 100 ng/mL KaryoMAX™ Colcemid™ Solution (Thermo Fisher Scientific, Cat. 15212012) for 2 hours. Cells were subsequently harvested using Versene Solution (Thermo Fisher Scientific, Cat. 15040066) and collected in PBS (Thermo Fisher Scientific, Cat. 10010023). Karyotype analysis was carried out at Sahlgrenska University Hospital (Gothenburg, Sweden).

### Flow Cytometry

Pluripotency of healthy control and patient iPSC lines was characterized by flow cytometric detection of SSEA4 and OCT4, together with SSEA1 used as a negative pluripotency control. Cells were stained using the BD Stemflow™ Human and Mouse Pluripotent Stem Cell Analysis Kit (BD, Cat. 560477) in accordance with the manufacturer’s protocol and analyzed on a BD FACSAria™ III flow cytometer.

### Bulk RNA Sequencing

Bulk RNA sequencing was used as an additional quality-control tool to characterize healthy control and patient iPSC lines and to support the characterization of NPCs and neurons. RNA sequencing was performed by BGI using the DNBSEQ platform, and gene expression data were provided by BGI as transcripts per million (TPM) values. For visualization, the provided TPM values of selected marker genes were log_2_(TPM + 1)-transformed and subsequently standardized using row-wise Z-score scaling. The resulting relative expression patterns across iPSCs, NPCs, and neurons were visualized as a heatmap.

### Stem Cell Differentiation to Cortical Neurons

Human iPS lines were differentiated to cortical glutamatergic neurons by following previously established protocol ^54^. iPSC lines were cultured in Biolaminin LN521 (Biolamina, Cat. LN521-05) coated plates in mTeSR+ medium (STEMCELL, Cat. 100-0276). Before starting differentiation, cells were cultured in specified coating agent and passaged with Versene (Thermo Fisher Scientific, 15040-066) for two weeks. After reaching 70-80% confluency, cells from two wells of six-well plates were transferred to one well to start differentiation in mTeSR+ medium with Y-27632 (10 µM; STEMCELL, Cat. 72302) to increase cell survival. During the first 10-12 days, cells were cultured with 10 μM SB431542 (Tocris, Cat. 1614/1) and 500 ng/mL Noggin (Bio-Techne R&D Systems, Cat. 3344-NG-050) by changing the media every day to induce neuroectodermal differentiation. Neural Maintenance Medium (NMM) was prepared fresh and contained the following: DMEM/F-12 with GlutaMAX (Thermo Fisher Scientific, Cat. 31331-028) and Neurobasal medium (Thermo Fisher Scientific, Cat. 12348-017) mixed at a 1:1 ratio, supplemented with insulin (2.5 µg/mL; Sigma-Aldrich, Cat. I9278), β-mercaptoethanol (55 µM; Thermo Fisher Scientific, Cat. 21985-023), MEM non-essential amino acids (0.5×; Thermo Fisher Scientific, Cat. 11140-035), sodium pyruvate (0.5 mM; Sigma-Aldrich, Cat. No. S8636), penicillin–streptomycin (25 U/mL penicillin and 25 µg/mL streptomycin; HyClone, Cat. SV30010), N-2 supplement (0.5×; Thermo Fisher Scientific, Cat. 17502-048), B-27 supplement (0.5×; Thermo Fisher Scientific, Cat. 17504-044), and additional GlutaMAX (1 mM; Thermo Fisher Scientific, Cat. 35050-038). From DIV10-12 to DIV25-26, the cells were passaged using Dispase (STEMCELL, Cat. 07913) every 2-3 days, depending on culture density and morphology, to enrich neural progenitor cells and reduce non-neural cells. The day after Dispase, the medium was changed with fresh NMM. For approximately 4 days following the initial Dispase passage, cells were cultured in NMM supplemented with FGF2 (20 ng/ml; Bio-Techne R&D Systems, Cat. 233-FB-025) to promote cell growth. Around DIV25-26, the first StemPro Accutase Cell Dissociation Reagent (Thermo Fisher Scientific, Cat. A1110501) passage was performed to obtain a single-cell suspension of neural progenitors, which were replated at approximately 70-80% confluency to support progenitor expansion. Cells were subsequently passaged using Accutase every 2-3 days at a 1:2 split ratio until final seeding. The final seeding was performed on DIV35. Cells were lifted using Accutase and seeded at a density of 50.000 cells/cm^2^ onto a plate coated with Poly-L-Ornithine (PLO, Merck Life Science, Cat. ABP4957-50ML) and Biolaminin (Biolamina, Cat. LN521-05). After final seeding, the medium was changed every second day using NMM. Brightfield images were acquired regularly using a Nikon Ts2-FL inverted fluorescence microscope (Nikon) to monitor cell morphology and differentiation throughout the culture period.

### PCR-targeted Long Read Sequencing of *CACNA1C* Transcripts

PCR-targeted long-read sequencing of full-length *CACNA1C* transcripts was performed on total RNA isolated from 32 samples across iPSC, NPC, and neuronal differentiation stages using a previously established protocol ^20, 55, 56^. Briefly, 400 ng of total RNA from each sample was reverse transcribed using Maxima H Minus Reverse Transcriptase (Thermo Scientific) with oligo(dT) priming according to the manufacturer’s instructions. To improve reverse transcription efficiency for GC-rich templates, the optional 65°C incubation step was included prior to cDNA synthesis. Reverse transcription reactions containing 20 U of enzyme were incubated at 50°C for 30 min.

Full-length *CACNA1C* amplicons were generated using a two-step PCR strategy to improve amplification efficiency of low-abundance transcripts and barcode-adapted products. The first PCR was performed using PrimeSTAR GXL DNA Polymerase (Takara) with cDNA equivalent to 100 ng starting RNA and primers listed in **Supplementary Table 9**. Amplification consisted of 25 cycles of 98°C for 10 s, 57°C for 15 s, and 68°C for 7 min, preceded by an initial denaturation at 98°C for 2 min and followed by a final extension at 68°C for 5 min. PCR products were purified using the Monarch PCR Cleanup Kit (NEB) and subsequently used as template for a second amplification consisting of 12 additional cycles under identical thermal conditions.

The resulting ∼6.5 kb amplicons were gel purified using 1% agarose/TAE gels stained with GelGreen (Biotium) and extracted using the Monarch DNA Gel Extraction Kit (NEB). Barcoding PCR was subsequently performed using PCR Barcoding Expansion 1–96 (EXP-PBC096, Oxford Nanopore Technologies) and PrimeSTAR GXL polymerase for 15 cycles of 98°C for 10 s, 62°C for 15 s, and 68°C for 7 min, followed by a final extension at 68°C for 5 min. Barcoded products were purified using the Monarch PCR Cleanup Kit (NEB).

DNA concentrations were quantified using the Qubit Broad Range DNA assay (Thermo Fisher Scientific), and samples were pooled in equimolar amounts. To remove low-molecular-weight DNA fragments, pooled libraries were purified using 0.4x Mag-Bind TotalPure NGS beads (Omega Bio-tek). Final library quality and fragment size distribution were assessed using Agilent Genomic DNA ScreenTape and the Qubit Broad Range DNA assay.

### Long-read processing and isoform identification

For sequencing, 1 µg of pooled library was prepared using the Ligation Sequencing Kit (SQK-LSK110, Oxford Nanopore Technologies) and sequenced on FLO-MIN106 flow cells for 72 h. Sequencing generated approximately 6.92 million reads. Base calling and barcode demultiplexing were performed using Guppy v6.4.8 (Oxford Nanopore Technologies).

Isoform discovery and quantification from nanopore long-read amplicon sequencing data was carried out using the IsoLamp pipeline v1.0.0 ^21^, (https://github.com/ClarkLaboratory/IsoLamp). Briefly, passed Nanopore reads with a *q*-score ³ 7, were down-sampled to 10,000 reads per barcode and mapped to the human reference genome (GENCODE v47) using Minimap2 ^57^. Aligned reads with an error-rate above 10% were removed. High-accuracy reads were used for isoform discovery with Bambu ^58^ using the parameters: novel discovery rate (NDR) = 1 and min.fractionByGene = 0.001. Isoform quantification was performed with Salmon ^59^ in alignment-based mode. Isoforms were then filtered to retain those that overlapped known primers coordinates and met an expression threshold of 500 TPM in a minimum of 8 (25% of total) samples. PCA plots and expression heatmaps were generated using the R statistical programming language v4.5.0 ^60^ and the packages ggplot2 ^61^ and ComplexHeatmap ^62^.

### Quantitative Polymerase Chain Reaction

Total RNA was isolated using the ReliaPrep™ RNA Miniprep Systems (Promega, Cat. Z103B). Whole-cell lysates collected in BL buffer supplemented with 1-thioglycerol to be lysed, mixed with isopropanol, washed with RNA wash solution, and incubated with DNase I incubation mix for 20 min. Samples were washed three times and collected in Nuclease-Free water in an elution tube by following the manufacturer’s instructions. The concentration of total RNA was measured via NanoDrop™ 2000/2000c Spectrophotometers. The quality of RNA was evaluated to A230/A260, A260/A280 values. Total RNA samples were converted to cDNA using a final concentration of 500 ng/µL RNA in RNAse-free water. cDNA was synthesized with a High-Capacity cDNA kit with RNase inhibitor (Applied Biosystems, Cat. 4387406) according to the manufacturer’s instructions. The cDNA was converted using the 2720 Thermal Cycler (Applied Biosystems) at 37°C for 60 min and 95°C for 5 min. Quantitative polymerase chain reaction (qPCR) was performed with a reaction volume of 15 μL containing the forward and reverse primer pair and Fast SYBR Green Mix (Thermo Fisher Scientific Europe, Cat. 4385612). The reactions were carried out in 96-well optical microtiter plates using the QuantStudio 3 Real-Time PCR System Software (Applied Biosystems). CT values for each targeted exon and *CACNA1C* exon 3 were averaged across two replicates and first normalized to *GAPDH* using the 2^−ΔCT^ method. To account for differences in total *CACNA1C* transcript abundance, the expression of each targeted alternative exon was subsequently normalized to exon 3, which was used as a constitutive exon representing total *CACNA1C* expression. Relative exon abundance was calculated as the ratio of *GAPDH*-normalized expression of the targeted exon to *GAPDH*-normalized expression of exon 3. The primers used in this study are provided in **Supplementary Table 9.**

### Cryopreservation

Dorsal forebrain organoids were fixed in 4% paraformaldehyde overnight at 4°C. The organoids were then washed with PBS -/- and transferred to 30% sucrose overnight at 4°C. Subsequently, excess sucrose was removed from the surface of the organoids, and the organoids were transferred into embedding molds containing embedding medium. The samples were frozen on dry ice and stored at −80°C until sectioning. For immunohistochemistry, 10 µm-thick sections were cut using a cryostat (Leica CM1950). Cryosections were collected onto SuperFrost Plus adhesion microscope slide (VWR, Cat. 631-0108) and stored at −20°C until immunostaining.

### Immunocytochemistry of 2D cultures

For 2D cultures, cells were seeded and cultured in 8-well ibidi plates (ibidi, Cat. 80826). Fixation was performed with 4% Paraformaldehyde (PFA) for 20 min at room temperature. The fixed cultures were washed three times for 5 min with 1X PBS -/- and permeabilized with 0.3% Triton-X in PBS -/- for 20 min. The fixed and permeabilized cultures were blocked with 2% BSA solution (Merck, Cat. A1470-100G) for 1 hour. Primary and secondary antibodies were prepared by following the manufacturer’s instructions. The fixated, permeabilized, and blocked cells were treated with primary antibody overnight at 4°C. After primary antibody incubation, the culture was washed 3 times for 5 minutes with PBS-/-. Secondary antibody incubation was performed for 1 hour at room temperature. After the secondary antibody incubation, the culture was washed 3 times for 5 minutes with PBS to remove unbound antibody.

### Immunofluorescence staining of organoid cryosections

Cryosections were washed with PBS -/- to remove residual OCT medium and then permeabilized with 0.3% Triton X-100 in PBS for 20 min. The sections were subsequently blocked with 2% BSA in PBS for 1 h at room temperature. Primary antibodies were diluted in blocking solution (2% BSA in PBS) and incubated with the sections overnight at 4°C. The following day, sections were washed three times with PBS for 5 min each to remove unbound primary antibodies. The sections were then incubated with the appropriate secondary antibodies, diluted in blocking solution, for 1 h at room temperature. DAPI was included during secondary antibody incubation at 1:1000 dilution. Following incubation, the sections were washed three times with PBS to remove unbound secondary antibodies. Finally, mounting medium was applied to the sections, and coverslips were placed over the samples prior to imaging.

### Confocal Imaging

Confocal imaging was performed using a Nikon Eclipse Ti-E confocal microscope equipped with 10x and 20x objectives. For organoid cryosections, images were acquired from randomly selected peripheral regions of each organoid to minimize the inclusion of the necrotic core. The primary and secondary antibodies used in this study, including their respective dilutions, are listed in **Supplementary Table 10.**

### Forebrain Dorsal Organoid Generation

Human ESC lines were cultured on Matrigel (Corning, Cat. 15505739) coated plates in mTeSR+ medium (STEMCELL, Cat. 100-0276). For the generation of dorsal forebrain organoids from ES lines, STEMdiff™ Dorsal Forebrain Organoid Differentiation Kits (STEMCELL, Cat. 08620) were utilized by following the protocols adapted from ^35^. Briefly, hESCs were dissociated into single cells using Accutase and seeded at a density of 3 million cells per AggreWell™ 800 24-well (STEMCELL, Cat. 34815) plate to form spheroids. The cells were cultured in the AggreWell™ 800 24-well (STEMCELL, Cat. 34815) plate system undisturbed for the first six days to initiate neural induction through SMAD pathway inhibition. Following this initial neuralization phase, spheroids were carefully transferred to ultra-low attachment plates and maintained in the forebrain expansion medium to promote regional patterning. Media were fully exchanged every other day. This phase continued until day 25 of differentiation. From day 25 to day 43, the cultures were transitioned to a forebrain organoid medium to provide neurotrophic support and further maturation. Full medium changes were consistently performed every other day throughout this period. For extended culture periods, the STEMdiff™ Neural Organoid Maintenance Kit (STEMCELL, Cat. 100-0120) was used to maintain organoid viability.

### Electrophysiology

#### Cloning

The splice variants *CACNA1C*-217 and *CACNA1C*-241 were generated using a seamless DNA assembly system. The *CACNA1C*-77 plasmid was used as the starting construct. Exon containing DNA fragments required to generate the targeted splice variants were synthesized as gBlocks (Integrated DNA Technologies, IDT). The target backbone was linearized by a touchdown PCR using Q5 High-Fidelity DNA Polymerase (New England Biolabs, Cat: M0491S). PCR-amplified backbone fragments were gel-purified using NucleoSpin™ Gel and PCR Clean-up Kit (Macherey-Nagel, Cat: 11992242). The purified backbones and the corresponding synthetic exon fragments were assembled using the NEBuilder HiFi DNA Assembly Mix (New England Biolabs, Cat. E2621) at a 1:2 vector-to-insert molar ratio and a total DNA amount of 0.2 pmol, according to manufacturer’s instructions. The sequences of the resulting plasmids constructs were verified by Sanger sequencing or Oxford Nanopore Technologies (ONT) sequencing.

#### Oocyte preparation

Defolliculated *Xenopus laevis* oocytes (stage V–VI) were either obtained commercially (Ecocyte; grade I) or prepared in-house as previously described ^63^. All procedures involving animals were approved by the Linköping Animal Care and Use Committee of the Swedish Board of Agriculture (permit no. 14515).

The following constructs were used: human *CACNA1C* (α_1C_; splice variant 217 and 241; both containing exon 7 full, 8, 9*, 15 full, 22, 41 short, −45, generated in-house for heterologous expression in *Xenopus* oocytes), rabbit *CACNA2D1* (α_2_δ-1; UniProt P13806),and rat *CACNB4* (β_4_; UniProt D4A055; a gift from Pierre Charnet). cRNA was synthesized *in vitro* using the mMESSAGE mMACHINE T7 kit (Thermo Fisher Scientific), assessed by spectrophotometry and gel electrophoresis, and stored at −80°C.

Oocytes were injected at the equator (Nanoliter 2020 injector, World Precision Instruments) with 50 nL of a cRNA mixture containing 0.3 μg/μL of α_1C_, α_2_δ-1, and β_4_. Following injection, oocytes were incubated for 3–4 days at 17 °C in a solution containing: 0.5× Leibovitz’s L-15 medium (Corning), diluted with H_2_O and supplemented with 1% heat-inactivated horse serum (Gibco), penicillin/streptomycin (10 U/L and 100 mg/L; Capricorn Scientific), and amikacin (100 mg/L; Fisher BioReagents).

#### Cut-open oocyte Vaseline gap (COVG) recordings

COVG is a low-noise, fast voltage-clamp technique ^64–66^. Before mounting in the recording chamber, each oocyte was injected with 100 nL of a solution containing 100 mM BAPTA•4K (Invitrogen) and 10 mM HEPES (pH 7.0) to suppress endogenous calcium- and barium-dependent chloride currents, as previously ^41, 67^. During activation experiments, the external solution contained (in mM): 120 Na-methanesulfonate (MeS), 10 K-MeS, 2 Ba(MeS) , and 10 HEPES (pH 7.0). Barium was used as a charge carrier to prevent calcium-dependent regulation. During inactivation experiments, Ca² current recordings were performed using the external solution contained (in mM): 120 Na-methanesulfonate (MeS), 10 K-MeS, 2 Ca(MeS) , and 10 HEPES (pH 7.0). For Ba² current recordings, Ca² was replaced by 10 mM Ba(MeS) , to produce the same level of charge screening. The internal solution consisted of (in mM): 120 K-glutamate and 10 HEPES (pH 7.0). The intracellular micropipette solution contained (in mM): 2700 Na-MeS, 10 NaCl, and 10 HEPES (pH 7.0).

A Clampator One CA-1B amplifier (Dagan Corporation) was operated in COVG mode. Current was low-pass filtered with cut-off frequency = 5 kHz using the onboard Bessel filter and acquired by a Digidata 1550B1 digitizer and pClamp 11.2.1 software (Molecular Devices) at 25 kHz.

#### Data analysis

The voltage dependence of activation was obtained from the peak tail current at −80 mV (*I*_tail_) and plotted against the test potential. Its voltage dependence was empirically characterized by fitting to the sum of two Boltzmann functions as:

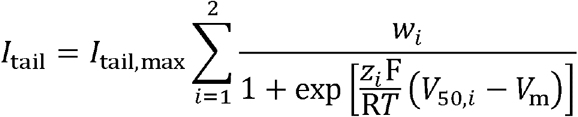

where *I*_tail,max_ was the maximal tail current, *w* the relative weight such that *w*_1_ + *w*_2_ = 1, *z* was the valence, F and R were the Faraday and gas constants, respectively, *T* was the absolute temperature, *V*_50_ was the half-activation potential and *V*_m_ was the test membrane potential. The effective *V*_50_ was calculated iteratively using *Solver* in Excel 365 (Microsoft).

The voltage dependence of inactivation was determined by fitting the fraction of non-inactivated current to a Boltzmann function:

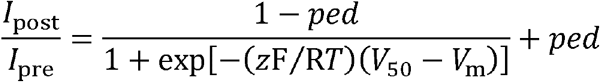

where *I*_pre_ and *I*_post_ represent current measured at 0 mV before and after the conditioning pulse, respectively, and *ped* corresponds to the non-inactivating pedestal current. Curve-fitting was performed by least squares using *Solver* in Excel 365.

#### Data Handling, Statistical Analysis, and Image Processing

Confocal immunofluorescence images were processed using NIS-Elements Advanced Research software (version 6.02.02) and Adobe Photoshop (version 27.3.0). Data visualization and statistical analyses were performed using GraphPad Prism (version 11.1.0; GraphPad Prism for macOS) and R (version 4.6.1). Nanopore sequencing data were statistically analyzed in R using the vegan package (version 2.7-6) for multivariate analyses, permute (version 0.9-10) for restricted permutations, and ggplot2 (version 4.0.3) for visualization. RT-qPCR data were analyzed and visualized using GraphPad Prism.

Overall, the *CACNA1C* isoform composition was analyzed using Hellinger–Euclidean distance based permutational multivariate analysis of variance (PERMANOVA) with 9,999 permutations. Bray-Curtis and robust Aitchison distances were used for sensitivity analyses. The primary analysis included a balanced repeated measures cohort of 30 samples from 10 donors at the iPSC, NPC, and neuronal stages. Permutations for developmental stages were restricted within donors to preserve repeated measure structure, whereas genotype, disease status, sex and age were evaluated in separate stage adjusted models using whole donor permutations. Pairwise developmental stage comparisons were performed using PERMANOVA with Holm correction. Multivariate dispersion was assessed using PERMDISP.

Developmental changes in reconstructed exon/splice features were evaluated using Friedman tests, with Benjamini-Hochberg (BH) correction. Pairwise developmental stage comparisons were performed using paired Wilcoxon signed-rank tests with Benjamini-Yekutieli (BY) correction. Between group comparisons of exon/splice feature contributions between patients and controls, rs1006737 A and G homozygotes, and females and males were performed separately within each developmental stage using Welch’s t-tests with Benjamini–Hochberg (BH) correction applied across features within each stage and group comparison. Total exon inclusion, including co-inclusion of exon pairs 21/22 and 31/32, was evaluated separately as an exploratory analysis using the same repeated-measures tests. Statistical significance was defined as an adjusted p < 0.05 within the corresponding testing family.

For exon abundance quantification in dorsal forebrain organoids done by RT-qPCR experiments, statistical differences in relative abundance across developmental stages were analyzed using two-way mixed-design analysis of variance (ANOVA), with developmental time as the between-subject factor and exon identity as the within-subject (repeated-measures) factor. Exon measurements obtained from the same organoid were treated as paired observations, whereas organoids collected at different times points were considered independent. Post hoc comparisons between exons at individual developmental stages were performed using Šídák’s multiple-comparisons test. Data are presented as mean ± SEM, and statistical significance was defined as *p* < 0.05.

The *p*-values for the differences between the voltage-dependence parameters of Ca_V_1.2-217 and Ca_V_1.2-241 were estimated using two-tailed, unpaired Student’s *t*-tests in Excel 365.

All figures were created using Adobe Illustrator, Adobe Photoshop, and open-source bioimaging tools (NIHBIOART) from the NIH along with Bioicons.

## Tables

**Supplementary Table 1**

Human iPS lines from the St. Göran Bipolar Project were used in the experiments shown in **Figure 1**.

**Supplementary Table 2**

Background information for bipolar patients and healthy controls from the St. Göran Bipolar Project, including sample ID, *CACNA1C* allele, age, sex, and bipolar subtype (NOS: not otherwise specified, N/A: not available).

**Supplementary Table 3**

Identity (Column A), relative abundance (Column B) and ranking (Column C) of the 60 *CACNA1C* isoforms identified in the present study. Column D shows whether the isoform has been reported in long-read RNA sequencing studies of human brain ^20, 21^. Column E shows the nucleotide sequence of each isoform.

**Supplementary Table 4**

Statistical analysis for association of *CACNA1C* isoform composition with developmental stage and donor characteristics. Worksheet 1-3 report Hellinger PERMANOVA (primary analysis), Bray-Curtis sensitivity, and robust Aitchison sensitivity. Worksheet 4-5 report PERMDISP and post-hoc stage comparisons. P_value and PERMDISP_P are raw P-values; P_Holm and PERMDISP_P_Holm are Holm-adjusted values. Significance follows Holm-adjusted *p* < 0.05; P_BH is provided for reference.

**Supplementary Table 5**

Exon annotation of all transcripts defined in Nanopore Sequencing. *CACNA1C* transcript models were visualized using the UCSC Genome Browser on the human GRCh38/hg38 genome assembly. Transcript structures, including both annotated and novel isoforms, were represented in BED format. Exon coordinates were derived from the BED block start positions and block sizes. Exon-level information was used to generate a binary transcript-by-exon matrix, in which 1 and 0 indicated the presence and absence of each exon, respectively.

**Supplementary Table 6**

Isoform-derived *CACNA1C* exon usage by developmental stages and donor characteristics. Worksheets report stage and stage-specific averages by diagnosis status, rs1006737 genotype and sex, respectively. Values were reconstructed from isoform abundances and expressed as percentages of the total *CACNA1C* isoform pool. It includes all available samples (n=12 iPSC, n=10 NPC, and n=10 Neuron).

**Supplementary Table 7**

Statistical results for *CACNA1C* exon-splicing features across developmental stages and donor groups. It includes overall stage associations, post-hoc stage comparisons, and within-stage comparisons by diagnosis, rs1006737 genotype, and sex. Exploratory total-inclusion results for exons 21, 22, 31, and 32 were also presented. Significance follows BH-adjusted *p* for overall stage tests and donor-group comparisons, and BY-adjusted *p* for post-hoc stage comparisons, at <0.05.

**Supplementary Table 8**

Activation and inactivation voltage dependence parameters for Ca_V_1.2-217 and -241 (**Figure 5**). Errors are SEM.

**Supplementary Table 9**

List of primers and oligos used in this study, including sequence information.

**Supplementary Table 10**

Information on primary and secondary antibodies, including host, dilution ratio, supplier, and catalog number details.

**Supplementary Table 11**

Genomic coordinates of *CACNA1C* exons based on the human reference genome GRCh38/hg38. The table includes chromosome location, exon start (5’ and 3’ end) positions.

## Data

### Data availability

The *CACNA1C* transcript isoforms identified by long-read sequencing are provided in BED format in **Supplementary Data 1** and can be visualized using UCSC Genome Browser. The raw and processed long-read and bulk RNA-sequencing data have been deposited in the NCBI Gene Expression Omnibus (GEO) and are publicly available as of the date of publication. Clinical data are not publicly available and may only be accessed subject to appropriate ethical approval and a data access agreement. Requests for access should be directed to the corresponding author, Erik Smedler.

### Supplementary Data 1

#### *CACNA1C* isoforms identified by Nanopore sequencing

Transcript isoforms are provided in BED format.

### Conflict of interest

The authors have no conflicts of interest to declare.

## Funding

This work was supported by the Swedish Research Council (ES: 2022-02185; AP: 2022-00574), the Knut and Alice Wallenberg Foundation (ES: Wallenberg Centre for Molecular and Translational Medicine at the University of Gothenburg and ES & AP: Program for Academic Leaders in Life Science collaboration grant, KAW 2022.0326), Åke Wibergs stiftelse (ES), Magnus Bergvalls stiftelse (ES), Jeanssons stiftelse (ES), Söderström-Königska (ES), Svenska Läkaresällskapet (ES), Åhléns stiftelse (ES), Stiftelsen Psykiatriska forskningsfonden (ES), Fredrik och Ingrid Thurings stiftelse (ES), Märta Lundqvist (ES), Längmanska Kulturfonden (ES), Göteborgs läkaresällskap (ES), Konrad och Helfrid Johansson (ES), and Bror Gadelius (MÇ). Hjärnfonden/The Swedish Brain Foundation (ES: FO2025-0065 and AP: FO2024-0299-HK-122 and FO2025-0364-HK-257). BD^2^: Breakthrough Discoveries for thriving with Bipolar Disorder (PJH, OA, NH, AM), National Institute for Health Research (NIHR) Oxford Health Biomedical Research Centre (NIHR203316; PJH, OA, NH, AM). The findings and conclusions are those of the authors and do not necessarily reflect the views of the NHS, the NIHR, the UK Department of Health. The St. Göran study was funded by grants from the Swedish Research Council (2022-01643), the Swedish foundation for Strategic Research (KF10-0039), Hjärnfonden/the Swedish Brain foundation (FO2025-0004-HK-212), and the Swedish Government under the LUA/ALF agreement (ALFGBG-1005343).

## Acknowledgements

We thank members of the Pantazis lab for useful discussions on the manuscript, and members of the Elinder, Larsson, Liin and Pantazis labs for expert *Xenopus laevis* oocyte preparation. We are grateful to Pierre Charnet for his generous gift of the Ca_V_-channel β_4_ cDNA. We thank Isar Nassiri for help with base calling.

## Author contributions

Conceptualization and study design: E.S., M.Ç.; Clinical data and iPSC lines: C.M.S., M.L.; iPSC work, neural differentiation and model characterization: M.Ç., M.N.; Long-read sequencing and transcript analysis: N.A.L.H., O.A.A., A.W.M., P.J.H.; Bulk RNA sequencing and analysis: M.Ç., M.N.; Electrophysiological experiments and analysis: A.P., S.S.Y.; Statistical analysis: M.Ç., O.A.A.; Supervision: E.S., A.P., A.W.M., P.J.H.; Funding acquisition: E.S., P.J.H., M.L.; Initial manuscript draft: E.S., M.Ç.; Manuscript review and editing: all authors.

## Use of generative AI

ChatGPT (OpenAI) was used to assist with language editing and wording. The authors reviewed and revised the text and take full responsibility for the content of the manuscript.

---

**Supplementary Figure 1.**
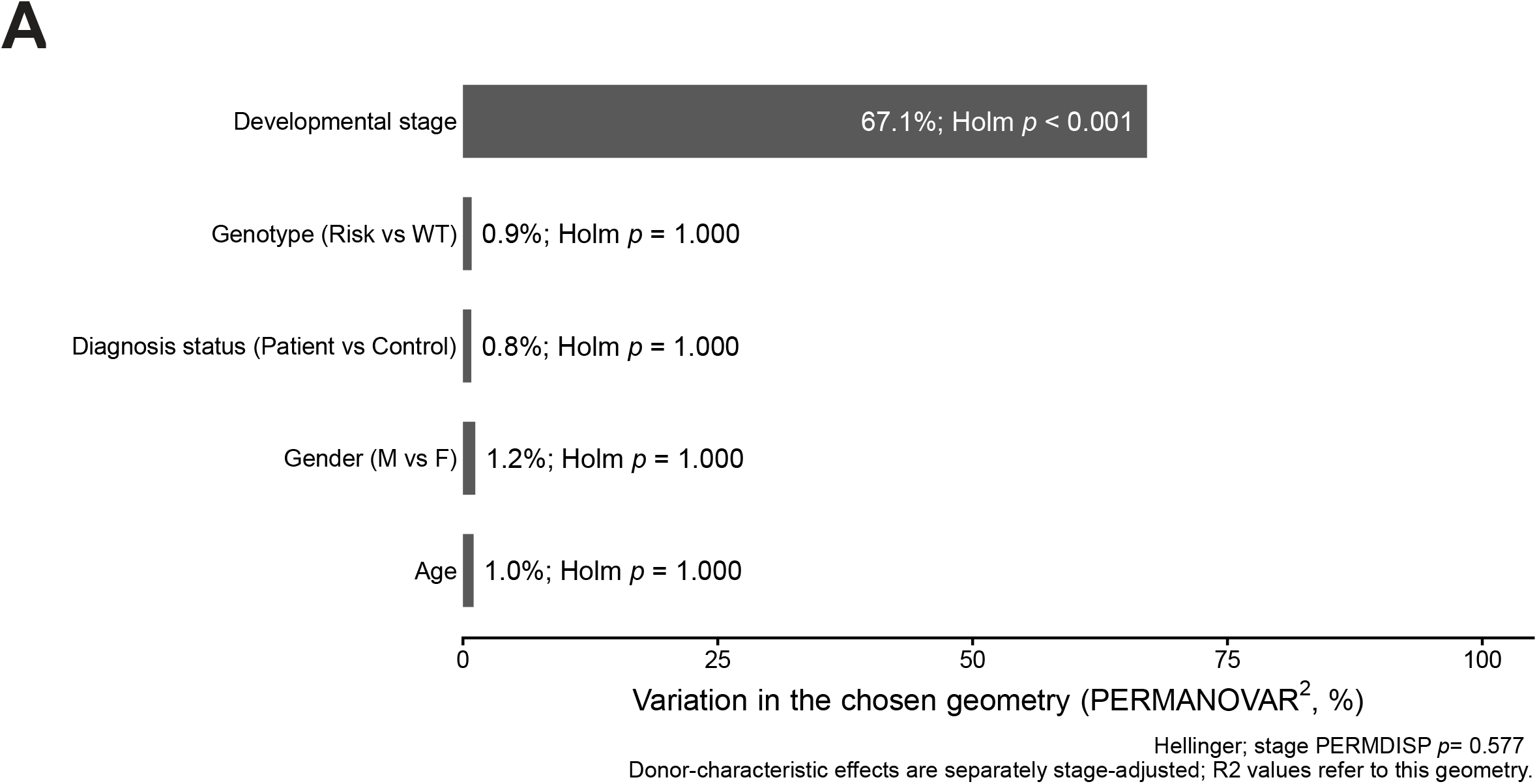
PERMANOVA analysis of parameters associated with *CACNA1C* isoform composition. **(A)** Bar graph representing the proportion of variation in *CACNA1C* isoform composition explained by developmental stage, rs1006737 genotype, diagnosis status, sex and age as determined by PERMANOVA based on Hellinger-Euclidean distances. Bars represent PERMANOVA R² values. Developmental stage explained the largest proportion of variation in isoform composition (R² = 0.671, *p*< 0.001), whereas rs1006737 genotype, diagnosis status, sex, and age showed substantially smaller effect sizes.

**Supplementary Figure 2.**
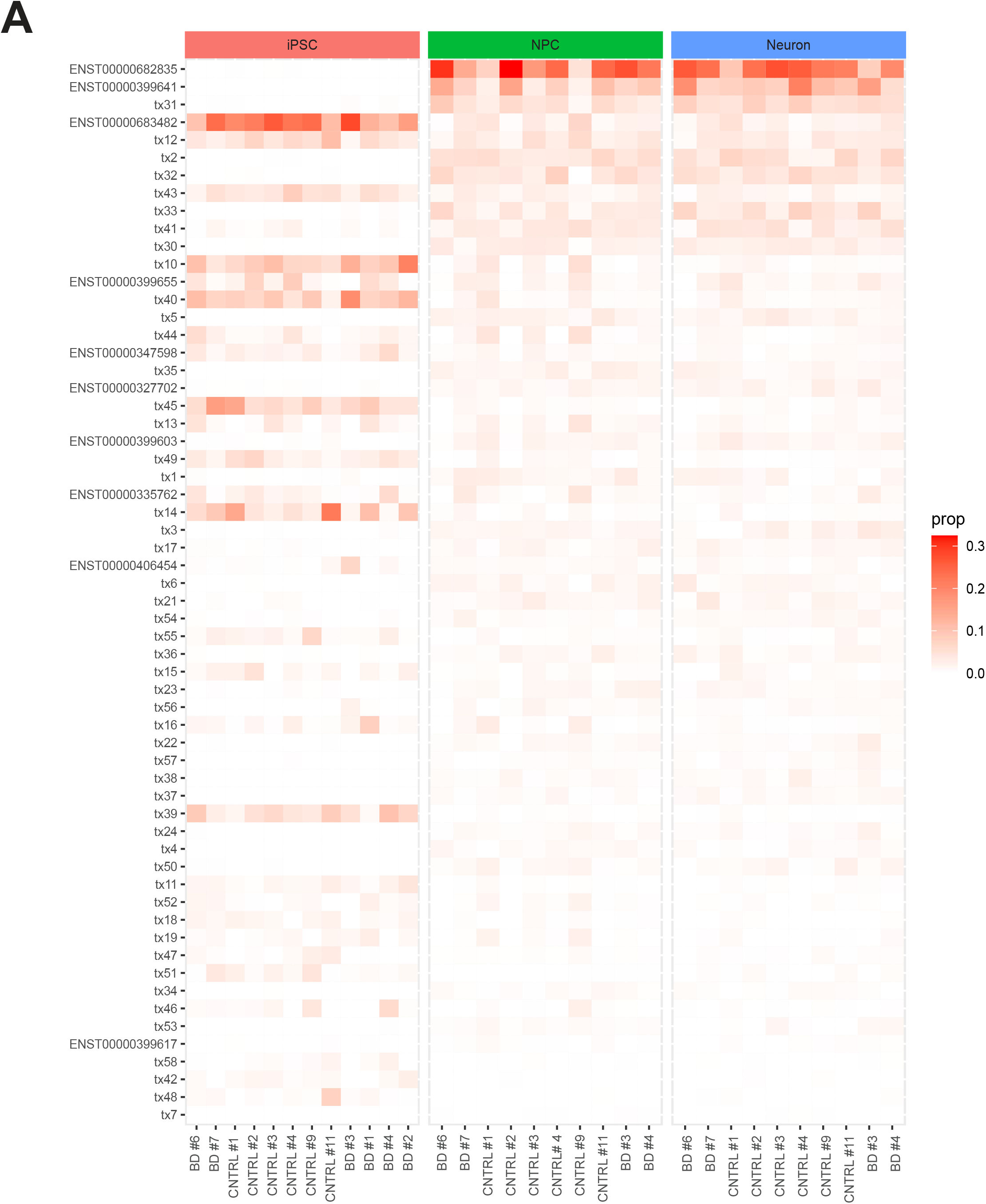
Detailed count proportions of different isoforms across all 32 samples. **(A)** Detailed table showing the count proportions of all 60 isoforms across the 32 samples analyzed by Nanopore sequencing. This table provides the detailed data corresponding to Figure 2C.

**Supplementary Figure 3.**
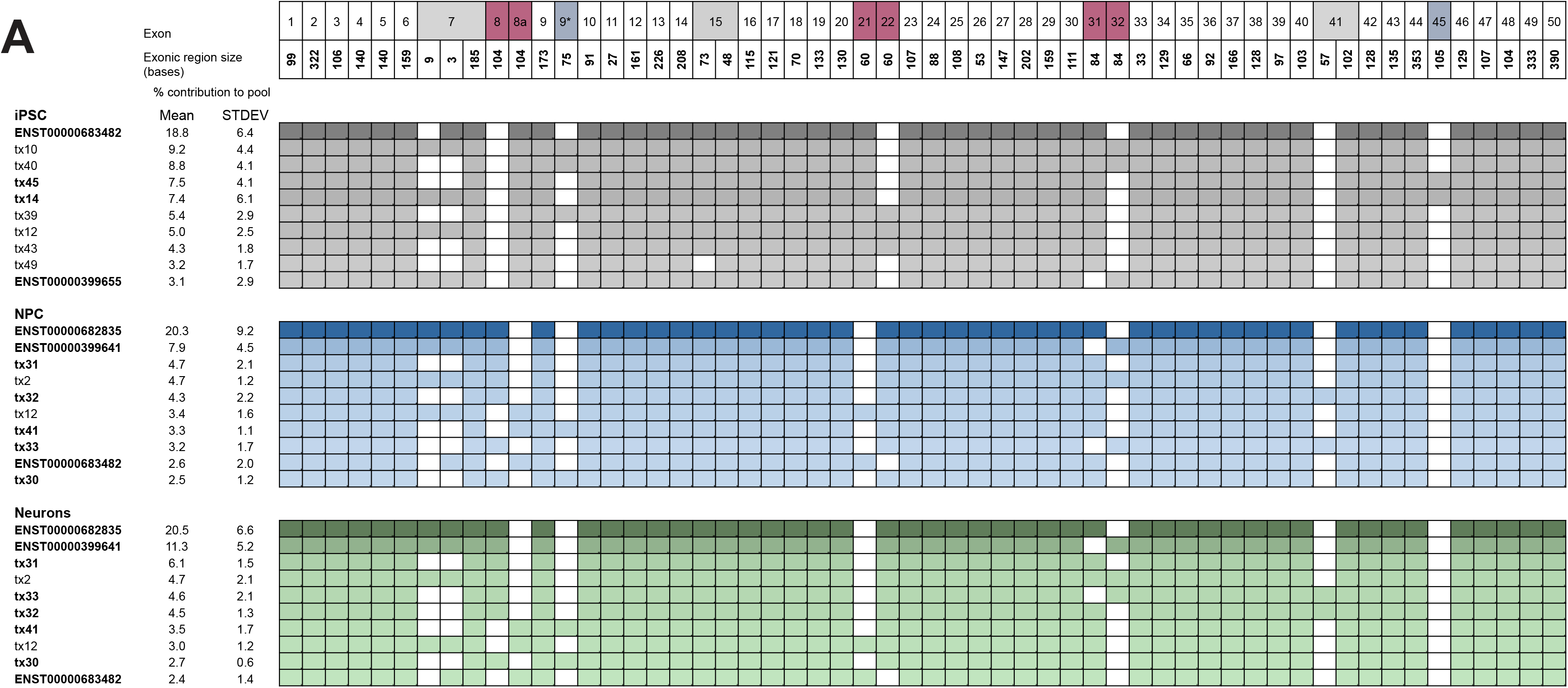
Exon composition and relative abundance of the ten most abundant *CACNA1C* isoforms identified across iPSC, NPC and neuronal stages. **(A)** Alternatively spliced exon regions are highlighted in gray (alternative 3’splice site), blue (casette exons), and dark pink (mutually exclusive exons). Exonic region sizes and exon structures are shown for the ten most abundant *CACNA1C* isoforms across stages, together with their percentage contributions to the transcript pool. Isoforms shown in bold are predicted to produce functional channels.

**Supplementary Figure 4.**
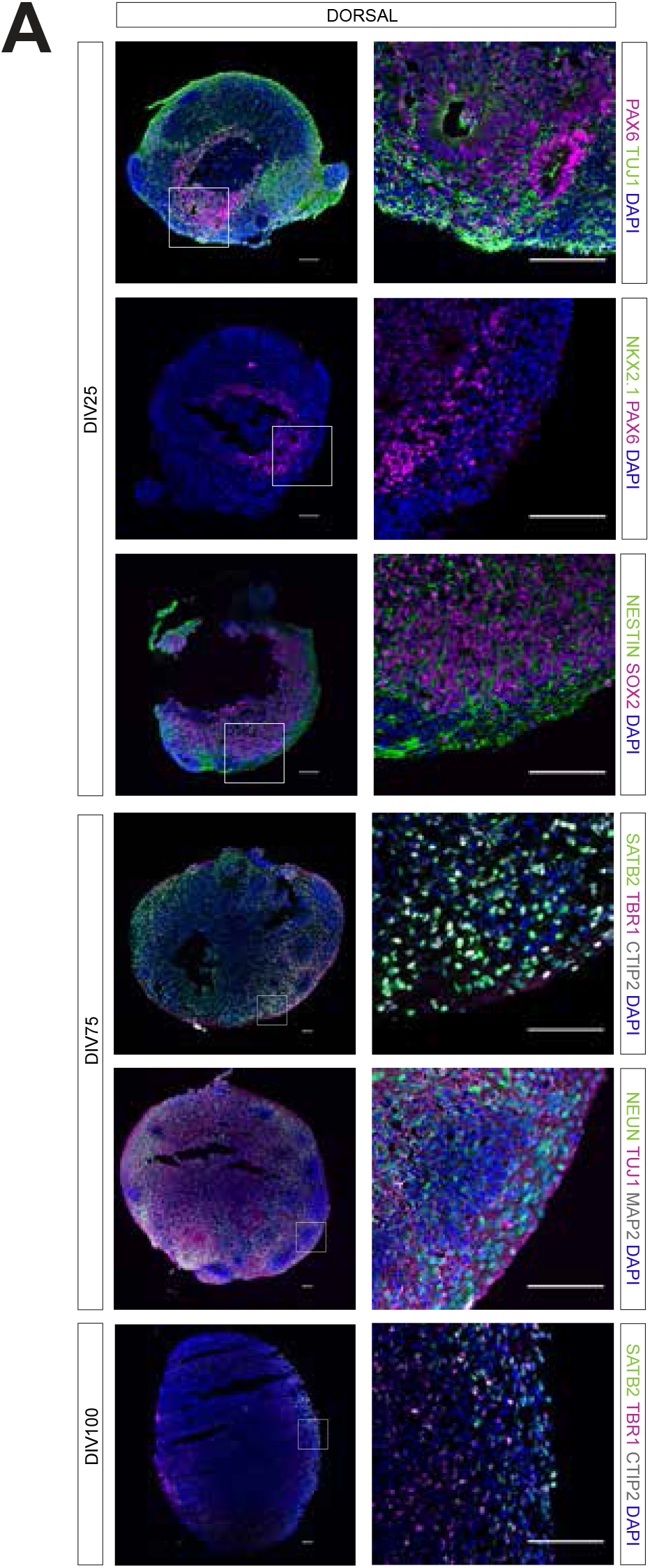
Validation of dorsal forebrain organoids across different time points. **(A)** Dorsal forebrain organoid identity and maturation were assessed also at DIV25, DIV75, and DIV100 using cortical layer markers (TBR1, CTIP2, SATB2), ventral markers (NKX2.1), neuronal maturation markers (TUJ1, NEUN, MAP2), progenitor marker (NESTIN, SOX2, PAX6). Representative immunostaining images were obtained from 10 µm sections of 1–3 independently processed organoids per developmental time point. Scale bar: 100 µm.

**Supplementary Figure 5.**
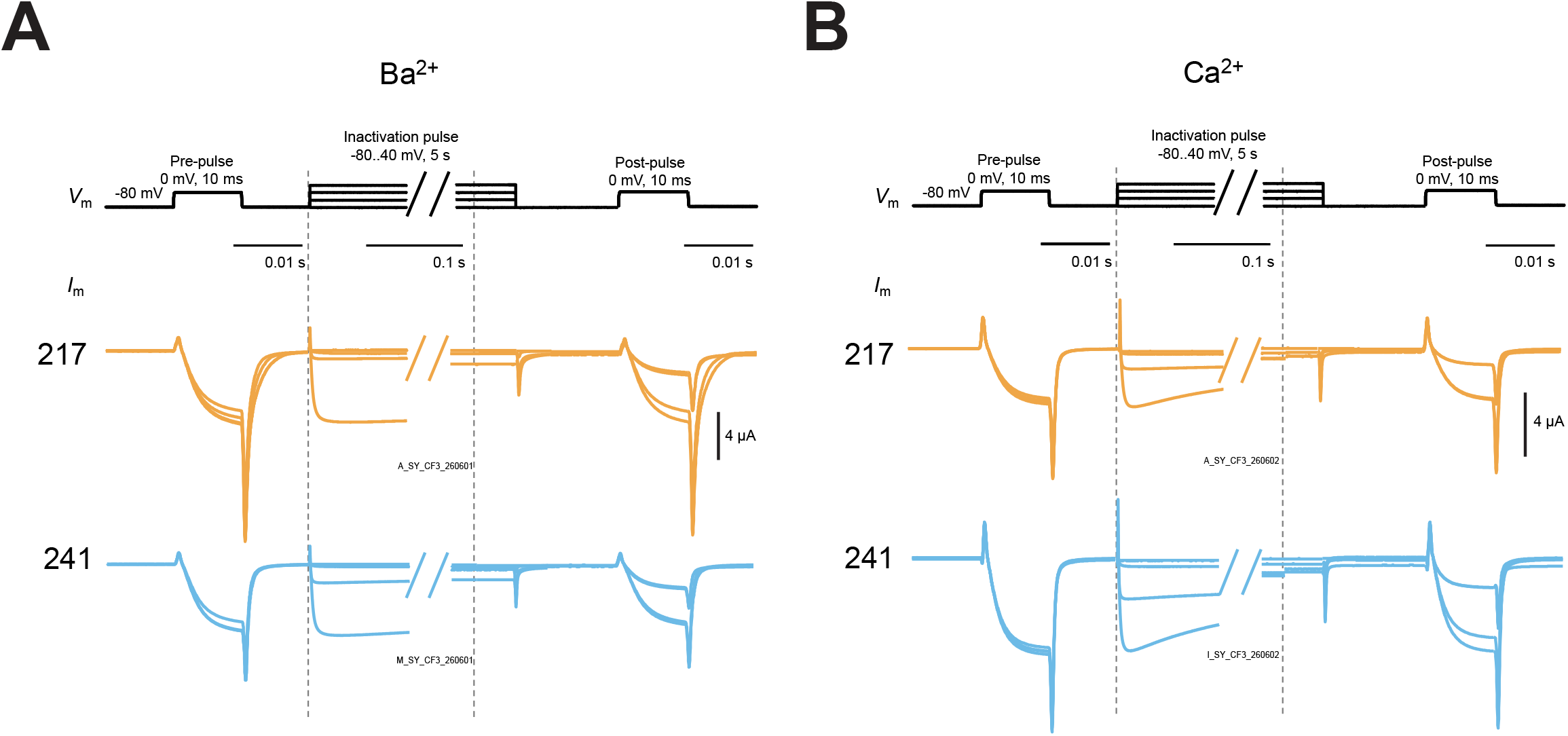
Representative inactivation traces. **(A)** Representative traces showing Ca_V_1.2-217 and Ca_V_1.2-241 inactivation in barium-containing external solution. Traces are shown normalized to the pre-pulse current in Figure 5E. **(B)** As in panel **A**, showing representative traces for inactivation in calcium-containing external solution. Traces are shown normalized to the pre-pulse current in Figure 5G.

## Notes

### Competing Interest Statement

The authors have declared no competing interest.

