## Supplementary Table 8 for "Developmentally-regulated splice remodeling shapes functional *CACNA1C* channel diversity during human neurodevelopment"

| Parameters | | 217 | 241 | $p$ |
|---|---|---|---|---|
| Activation | $V_{50,1}$ (mV) | −12 ± 2.6 | −13 ± 3.0 | 0.69 |
| | $z_1$ ($e_0$) | 2.7 ± 0.22 | 2.4 ± 0.17 | 0.32 |
| | $w_1$ | 0.59 ± 0.081 | 0.67 ± 0.077 | 0.51 |
| | $V_{50,2}$ (mV) | 25 ± 1.2 | 26 ± 1.9 | 0.59 |
| | $z_2$ ($e_0$) | 1.7 ± 0.31 | 2.1 ± 0.63 | 0.54 |
| | $V_{50}$, effective (mV) | −2.1 ± 3.9 | −4.5 ± 1.2 | 0.63 |
| | *n* (cells) | 4 | 3 | |
| Inactivation ($Ba^{2+}$) | $V_{50}$ (mV) | −25 ± 1.2 | −31 ± 1.5 | 0.016 |
| | $z$ ($e_0$) | 2.8 ± 0.20 | 2.8 ± 0.25 | 0.99 |
| | *ped* (%) | 26 ± 2.7 | 26 ± 1.4 | 0.86 |
| | *n* (cells) | 5 | 8 | |
| Inactivation ($Ca^{2+}$) | $V_{50}$ (mV) | −25 ± 1.2 | −31 ± 0.34 | 0.0031 |
| | $z$ ($e_0$) | 3.5 ± 0.23 | 3.7 ± 0.26 | 0.54 |
| | *ped* (%) | 23 ± 1.8 | 17 ± 3.0 | 0.11 |
| | *n* (cells) | 10 | 5 | |
